# Electrical synapses mediate visual approach behavior

**DOI:** 10.1101/2025.10.14.682373

**Authors:** Giovanni Frighetto, Mark Dombrovski, Lesly M. Palacios Castillo, Pratap Meera, Parmis S. Mirshahidi, Pegah S. Mirshahidi, Piero Sanfilippo, Andrea Vaccari, Pratyush Kandimalla, Volker Hartenstein, Yerbol Z. Kurmangaliyev, S. Lawrence Zipursky, Mark A. Frye

## Abstract

Detecting salient visual objects and orienting toward them are commonplace tasks for animals, yet the underlying neural circuits remain poorly understood. The fruit fly is an ideal model for a comprehensive analysis of feature detection mechanisms given its complete synaptic wiring diagrams, robust behavioral assays, and cell-type-specific gene expression datasets. We previously showed that columnar T3 neurons are required for saccadic orientation toward landscape features during flight. Here, we examine how signals downstream of T3 are processed in the central brain. We identify LC17 visual projection neurons as key postsynaptic targets: they receive strong excitatory input from T3, project to premotor brain regions, and are thus positioned to support visual approach. Using *in vivo* optical physiology and virtual reality behavior, we demonstrate that LC17 neurons are indeed necessary for object tracking during flight. Furthermore, we find that electrical synapses in LC17 are also required for tracking behavior. We show that the innexin Shaking B (*shakB*) is highly expressed in LC17 and localized to its dendrites, and genetic perturbations confirm its essential role for electrical coupling in this circuit. Our findings reveal mechanisms underlying visual approach, and highlight the interplay between electrical and chemical neurotransmission for rapid object detection and action selection.

## Main

Approaching or avoiding visual objects is a fundamental skill that animals need to safely explore their environment and exploit its resources. While the neural circuits underlying visual avoidance responses have been well studied in multiple model organisms including fruit flies^1–8^, zebrafish^9–12^, and mice^13–16^, the properties of visual approach behavior have only recently begun to be identified in vertebrates^17–20^ and the underlying circuit mechanisms remain largely elusive. The fruit fly is a genetically tractable model that exhibits complex visual behaviors and currently offers complete visual system chemical synaptic connectomes^21–23^, single-cell transcriptomes^24–26^, and a rich genetic toolkit for molecular and functional circuit analysis. To date, circuits underlying the control of approach behavior have been investigated by merely a handful of studies almost exclusively in walking flies^27–30^. Here, we focused on a visual pathway involved in encoding visual features for object tracking under the high-performance demands of visual flight control, thereby underscoring structure-function relationships. To clarify terminology, we use “approach” and “orient” interchangeably, reflecting the spatial dynamics of virtual reality flight in which a fly can control its orientation but cannot actually approach^31,32^. In a natural setting, orientation is *sine qua non* for approach, but it can also be a stand-alone behavior. To enable orientation, a visual object needs to be discriminated from the background by relying on features such as color, luminance, contours, texture, or relative motion^33^. Although the elementary direction-detecting neurons, T4 and T5, are an important component of motion vision^34^, their contribution to feature detection – and the role of their downstream partners in the lobula plate neuropil in object recognition – appears to be limited^27,35^. Many visual features are selectively processed by different visual projection neurons (VPNs) innervating the lobula neuropil^36–38^. Specifically, lobula columnar (LC) neurons^39^ – a class of more than 40 VPN types that maintain a dendritic retinotopic organization and a glomerular organization of axon terminals^22^ – show highly selective receptive field (RF) properties^38,40–42^. Different LC neuronal types are selective for features such as object shape, size, orientation, motion dynamics, and relay this information to type-specific glomeruli in the central brain^4,38,41–48^. In some cases, LC neurons connect directly to descending neurons (DNs) that command motor programs such as takeoffs^8,21,49^. In others, the circuits downstream of LC neurons are polysynaptic, involving multiple interneurons before sending information to DNs. One behavior that might require such a complex circuit organization is landing^50^. In this context, flying flies must segregate local landscape features from the distant visual background, select one, and track its position over time to approach and alight upon it. Flies are thus strongly attracted to vertical bars, likely because they resemble plant stalks or gaps in landscape foliage, and they orient and pursue these objects using a strategy of straight flight segments interspersed with rapid turns called body saccades^31,51,52^. Previously, we identified a type of numerous T-shaped neurons^53^, T3, which selectively supports saccadic bar tracking behavior^35^. These columnar cholinergic neurons are structurally and functionally connected with LC11^54,55^, a highly specialized feature detector tuned to small moving dark objects^43^. However, silencing and activation studies have highlighted a role of LC11 in freezing but not in visual approach behavior^43,55,56^. We therefore searched for additional downstream partners of T3 using two different connectomes of the optic lobe^21,22^. We identified LC17 neurons with dendrites in layer 2 and 3 of the lobula and axons in the posterior ventrolateral protocerebrum (PVLP) and here we investigated their role in visual approach using physiological, behavioral, anatomical, and molecular experiments.

## Results

### LC17 neurons are postsynaptic partners of T3

To identify downstream neurons receiving synaptic input from T3 neurons, we explored the female adult fly brain (FAFB) connectome^57^ which has been reconstructed, proofread, and annotated by the FlyWire community^21,58,59^. T3 neurons send most of their outputs to LC11 and LC17 neurons (Fig. 1a), which we confirmed using the more recent male optic lobe (MAOL) connectome^22^ (Fig. 1b). Within one of the bilateral optic lobes, ∼900 columnar T3 neurons connect to ∼170 LC17 neurons that cover the entire visual field (Fig. 1c) such that each LC17 receives input from about 20 individual T3 neurons (Fig. 1d). We next demonstrated T3-LC17 functional connectivity using optogenetic stimulation and calcium imaging^60,61^. This was performed on a two-photon excitation microscope equipped with a 3D-scanless holographic photoexcitation system (3D-SHOT^62^), allowing selective stimulation of discrete depth-restricted regions (∼10 μm in diameter) at T3 presynaptic terminals while recording from LC17 dendrites (Fig. 1e). First, we photostimulated a region covering approximately the size of two single T3 presynaptic terminals by pulsing light (50 Hz) at different power intensities and recorded the calcium response from the most proximal dendritic branch of an LC17 neuron (Fig. 1e,f). This provoked peak calcium responses in LC17 that increased with the power delivered (Fig. 1g). Next, after having identified the optimal power intensity (Extended Data Fig. 1a-c), we stimulated a sequence of three spatially separated holograms targeting T3 presynaptic terminals and recorded from a single LC17 dendritic compartment that opposed to the first stimulation site (Fig. 1e). This spatial pattern of photoactivation simulated an object moving back-to-front across the fly eye. The calcium response at one recording site indicated temporal integration across stimulations as well as spatial integration from input at distant sites, possibly due to dendro-dendritic interactions (Fig. 1h and Extended Data Fig. 1d,e). Curiously, imaging LC17 axons in response to different stimulation sequences of two directional holograms (front-to-back and back-to-front) revealed a subtle slow progressive increase in calcium accumulation to back-to-front motion (Extended Data Fig. 1f-h). This directional preference could be the result of a posterior-anterior gradient in the number of T3 cells providing inputs to LC17 neurons (Extended Data Fig. 1i). Finally, we took advantage of the slow kinetics that characterize the channelrhodopsin we used (Chrimson^60^) to modulate the efficiency of our photoactivations. By stimulating the neurons with the same optimal power, but constant light, the calcium responses did not accumulate and instead showed a decay with distance (Fig. 1i). Moreover, the stimulation of the three sites simultaneously provoked a response that resulted in a sub-linear summation of the responses to singular sites individually, which suggests neural inhibition (Fig. 1i).

**Figure 1.**
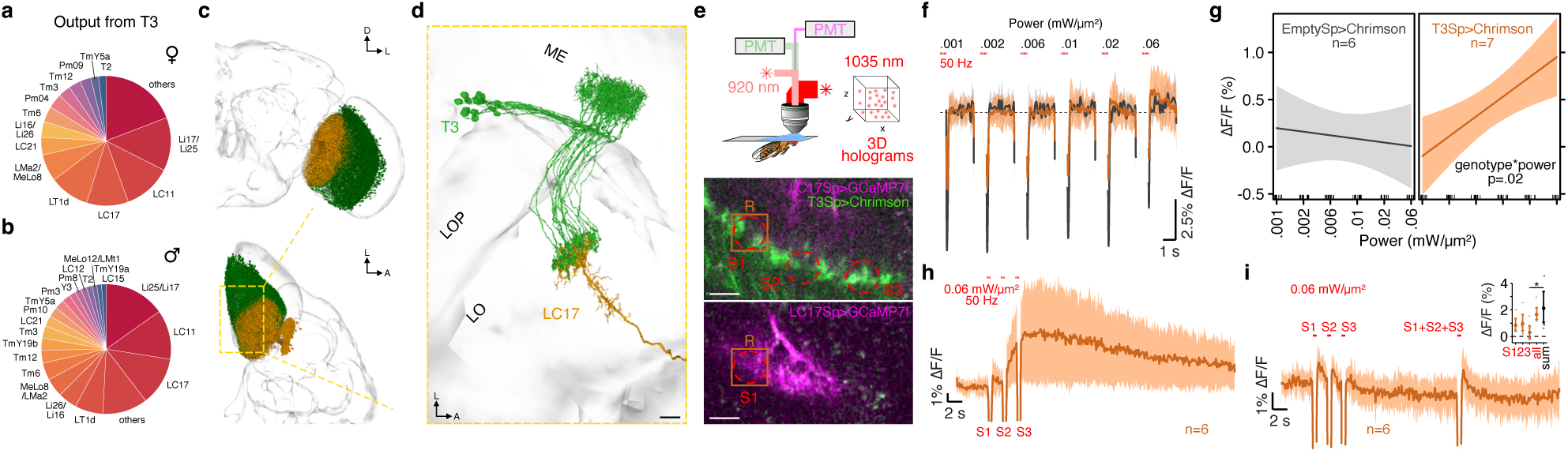
LC17 neurons are structurally and functionally connected to T3 neurons. **a**, Pie chart of the output neurons from T3 based on number of synapses in the female adult fly connectome (FAFB). **b**, Pie chart of the output neurons from T3, as in a, in the male adult fly optic lobe connectome (MAOL). **c**, Meshes of the FAFB brain with the whole population of LC17 (orange) and T3 (green) reconstructed neurons. Top: frontal view of the left optic lobe. Bottom: dorsal view of the left optic lobe. D, dorsal; L, lateral; A, anterior. **d**, Dorsal close-up of a single LC17 dendrites (orange) with its relative upstream T3 neurons (green). On average, ∼23 T3 cells send input to a single LC17 neuron. ME, medulla; LO, lobula; LOP, lobula plate. Reconstructed neurons represented as meshes from FAFB connectome. Scale bar, 10 µm. **e**, Top: schematic depicting the in vivo preparation used to stimulate T3 neurons with the 3D-SHOT method while recording calcium activity in LC17 neurons. PMT, photomultiplier tube. Center: two-photon image of LC17 dendrites expressing GCaMP7f (purple) and T3 presynaptic terminals expressing Chrimson tagged with tdTomato (green). Red dashed circles represent the 3D holograms stimulated (S1-3). Orange solid square (R) represents the region of interest (ROI) from which we recorded the calcium activity. Scale bars, 10 µm. Bottom: two-photon image of LC17 dendrites expressing GCaMP7f (purple) without the expression of Chrimson in T3 neurons. Single 3D hologram stimulated (S1) and ROI recorded (R) for the power sensitivity experiment. **f**, Power sensitivity experiment performed by recording calcium response (mean ± s.e.m.) in LC17 dendrite to a single 3D hologram stimulated with increasing power in flies expressing Chrimson in T3 neurons (orange; n = 7) and in control flies not expressing Chrimson (gray; n = 6). Each stimulation lasted 0.5 s and the light was pulsed at 50 Hz (single pulse duration, 5 ms) with an inter-stimulus interval (ISI) of 3 s. The dip in the traces at the beginning and at the end of each recording is an artifact generated by the PMTs shutting off during the stimulation period to prevent damage. **g**, Linear regressions fitting the average peak responses (200 ms window after the stimulation) across powers in Chrimson and no Chrimson expressing flies as shown in **f**. Two-way repeated measure ANOVA detected an interaction effect between genotype (Chrimson and no Chrimson) and ramping power (F_(1, 74)_ = 6.04, p=0.02). Shaded region represents confidence interval (C.I.) at 95%. **h**, Calcium response (mean ± s.e.m.) recorded from a single ROI placed over a LC17 dendrite during sequential stimulation of three holograms (S1-3) targeting T3 presynaptic terminals with increasing distance from the ROI (n = 6). Stimulation duration, 0.5 s; Stimulation frequency 50 Hz (5 ms single pulse); ISI, 2 s; Power, 0.06 mW/µm^2^. **i**, As in **h**, calcium response (mean ± s.e.m.) to sequential stimulation of three holograms followed by a final stimulation of the three holograms simultaneously (n = 6). Stimulation duration, 0.5 s; ISI, 2 s; Power, 0.06 mW/µm^2^. Inset: average peak responses (200 ms window after the stimulation) to the four stimulation conditions (orange). Small opaque dots represent single average peak responses. Bigger solid dots represent the sample mean and error bars ± 95% C.I. The black dot with error bar represents an artificial condition where the three sequential stimulations are summed together. Statistical analyses were performed using unpaired t-test corrected with Holm-Bonferroni method (*P<0.05).

To validate the physiological range of the photostimulation results, we visually presented the flies with bars moving in different directions. The visual stimuli elicited calcium signals comparable to the focal photoactivation of T3 neurons (Extended Data Fig. 1j), demonstrating that our optogenetic approach evoked robust responses that resemble the ones physiologically generated by the retinotopic activation of the input columnar neurons. Together, our all-optical interrogations^63^ align with connectomics data^58^: LC17 neurons are *bona fide* postsynaptic partners of T3. Furthermore, we show that sparse spatiotemporally patterned T3 activation evokes complex dendritic computations in LC17 including temporal integration and directional facilitation. This functional complexity might be the reason for the discrepancy between previous LC17 optogenetic stimulation experiments in freely walking flies, which triggered fast turning responses^38,64^, and our experiments in rigidly tethered flying and walking flies, which display a converse if not ambiguous phenotype (Extended Data Fig. 2a-c). Perhaps the behavior elicited by LC17 requires stimuli that comprise specific dynamic conditions (i.e., temporal and spatial patterning) or activation sequences that were achieved in freely moving flies but not in restrained preparations where ensembles of neurons are activated simultaneously.

### LC17 tuning properties corroborate a role in object tracking behavior

Saccadic bar tracking behavior in flight requires T3 neurons^35^. Among the top five T3 postsynaptic targets, LC11 and LC17 neurons share not only T3 but also T2a neurons as presynaptic partners (Extended Data Fig. 1k), making the majority of their synaptic input profile broadly similar. Earlier work established LC11 as small-object detectors, whose responses decrease with increasing stimulus height or width^43,55^. This flanking inhibition makes LC11 poorly tuned to vertical bars that strongly attract flies. Perhaps instead LC17 neurons integrate local T3 inputs to respond to bar-like stimuli^35^. To test this, we characterized LC17 dendritic visual responses using two-photon calcium imaging (Fig. 2a). We compared calcium activity of LC11 and LC17 neurons in response to vertical bars of different contrast polarities, as well as a motion-defined bar, and gratings of varying spatial wavelengths (Fig. 2b). Unlike LC11, LC17 showed robust responses to vertical bars of either ON or OFF contrast polarity (Fig. 2c and Extended Data Fig. 3a,b). Neither LC types responded to gratings, indicating active wide-field inhibition in LC17. Notably, the response amplitude of LC17 decreased as bar speed increased (Fig. 2d,e), mimicking T3 properties and supporting an integrate-and-fire conceptual model for triggering body saccades during bar tracking^31,35^. These findings align with previous studies that classified LC11 as small-object detectors, and LC17 as bar detectors with additional sensitivity to looming stimuli^46^. Yet, our earlier work reported difficulties detecting visually evoked calcium activity in LC17^41^. Here, three repetitions of the same moving bar revealed progressive response attenuation in LC17, suggesting strong habituation (Fig. 2f), and explaining the prior low-amplitude responses as the outcome of averaging over trials. LC11, by contrast, maintained weak but stable trial-by-trial response amplitude (Fig. 2f,g and Extended Data Fig. 3c): responses looked almost facilitated at bar speeds (i.e., 90-110 deg/s) near their tuning peak^54^ (Fig. 2f). Thus, habituation may regulate LC17 excitability and obscure averaged responses in long visual experiments^41,42^. Another recent study categorized LC17 exclusively as looming detectors^42^. To probe this, we compared LC17 to another VPN, the looming specialist LPLC2^4^. In response to RF-centered edges moved in 24 directions^65^, both LC17 and LPLC2 showed omni-directional sensitivity, with LC17 being slightly more biased for upwards motion (Fig. 2h). However, the response to looming was significantly stronger in LPLC2 (Fig. 2i). This indicates that LC17 looming responses originate from edge motion sensitivity rather than the radial expansion tuning seen in LPLC2 neurons (i.e., strong integration of motion from different cardinal directions)^4^. These results reveal that LC17 neurons exhibit strong edge detection, ON-OFF selectivity (i.e., full wave rectification), and omni-directional motion tuning. These properties make LC17 well suited to act downstream of T3 in supporting bar tracking behavior.

**Figure 2.**
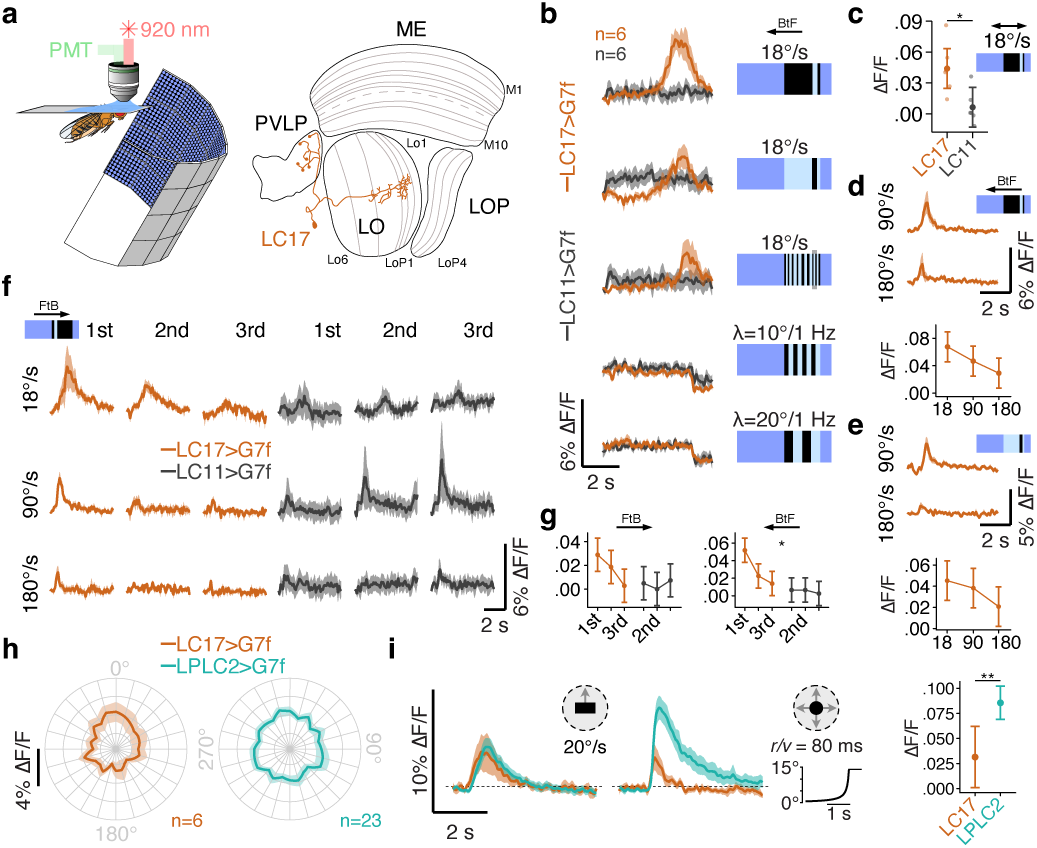
LC17 visual properties are distinguishable from small-object and looming detectors. **a**, Left: schematic of a head-fixed fly for two-photon calcium imaging while presented with visual stimuli from a surrounding LED display. Right: fly optic lobe neuropils (ME, medulla; LO, lobula; LOP, lobula plate; PVLP, posterior ventrolateral protocerebrum) with a LC17 neuron highlighted in orange. **b**, Calcium imaging responses in LC17 (orange) and LC11 (gray) neurons to a battery of different visual stimuli moving back-to-front (BtF) depicted on the right (n = 6 per group; mean ± s.e.m.). **c**, Maximum peak response in LC17 (orange) and LC11 (gray) for a moving ON bar (9° x 72°, width x height) at 18°/s (mean ± 95% C.I.). Small opaque dots represent single flies. **d**, Top: average responses (mean ± s.e.m.) to a BtF moving ON bar at two different velocities in LC17. Bottom: maximum peak responses at three different velocities (mean ± 95% C.I.). **e**, Same as in **d** for a moving OFF bar. **f**, Average calcium traces in response to an ON bar moving front-to-back (FtB) at different velocities over three repetitions in LC17 and LC11 (n = 6 per group, mean ± s.e.m.). **g**, Maximum peak response in LC17 (orange) and LC11 (gray) for FtB (left) and BtF (right) moving ON bar at 18°/s over three repetitions. **h**, Polar plots of the average peak responses in LC17 (orange; n = 6) and LPLC2 (turquoise; n = 23) to dark edges moving in 24 orientations (0°-345°) presented in a visual region above the fly eye’s equator (2 trials, mean ± s.e.m.). **i**, Left: average calcium response to an upward moving dark edge (left) and to a dark looming in LC17 (orange) and LPLC2 (turquoise) presented above the fly eye’s equator (mean ± s.e.m.). Right: maximum peak response in LC17 and LPLC2 to a dark looming (mean ± 95% C.I.). Data on LPLC2 are replotted from Dombrovski et al., 2025. Statistical analyses were performed using unpaired two-sided t-test (*P<0.05, **P<0.01, ***P<0.001).

### Gap junctions in LC17 support bar tracking behavior

Displacement of a motion-defined bar resembles the movement dynamics of flying by a nearby tree observed against the distant stationary visual surroundings. This stimulus effectively elicits approach via body saccades^31^. To reduce stimulus variables, the broadband spatial-frequency content of the moving bar matches that of the static background and is therefore discriminable only by its relative motion. We previously showed that T3 neurons are high spatial-frequency speed detectors and are fundamental for triggering bar-directed body saccades^35^. We next asked if postsynaptic LC17 neurons participated in the same visual control pathway. We used a magnetic-tether setup, wherein flies rotate freely in the yaw plane inside a cylindrical visual display, evoking clear body saccades (Fig. 3a)^66,67^. A motion-defined bar revolved around the fly while flight heading was videorecorded in flies with different neuron populations silenced (Fig. 3b). Silencing (i.e., hyperpolarizing) LC17 neurons by expressing an inwardly rectifying potassium channel (Kir2.1^68^) reduced bar tracking to a similar extent as silencing T3 neurons^35^ (Fig. 3c,d and Extended Data Fig. 4a), accompanied by a sharp drop in saccade frequency (Fig. 3e). Importantly, the effect of this manipulation was specific to the T3-LC17 circuit (i.e., silencing T2a neurons had no impact on tracking performance) and was specific to bar tracking. As predicted, the wide-field gaze stabilization (also known as optomotor response) supported by direction-detecting T4/T5 neurons^69^ was preserved in LC17-silenced flies, but compromised in T4/T5-silenced flies (Extended Data Fig. 4b). Thus, LC17 is a key peripheral component of the T3 pathway for bar discrimination.

**Figure 3.**
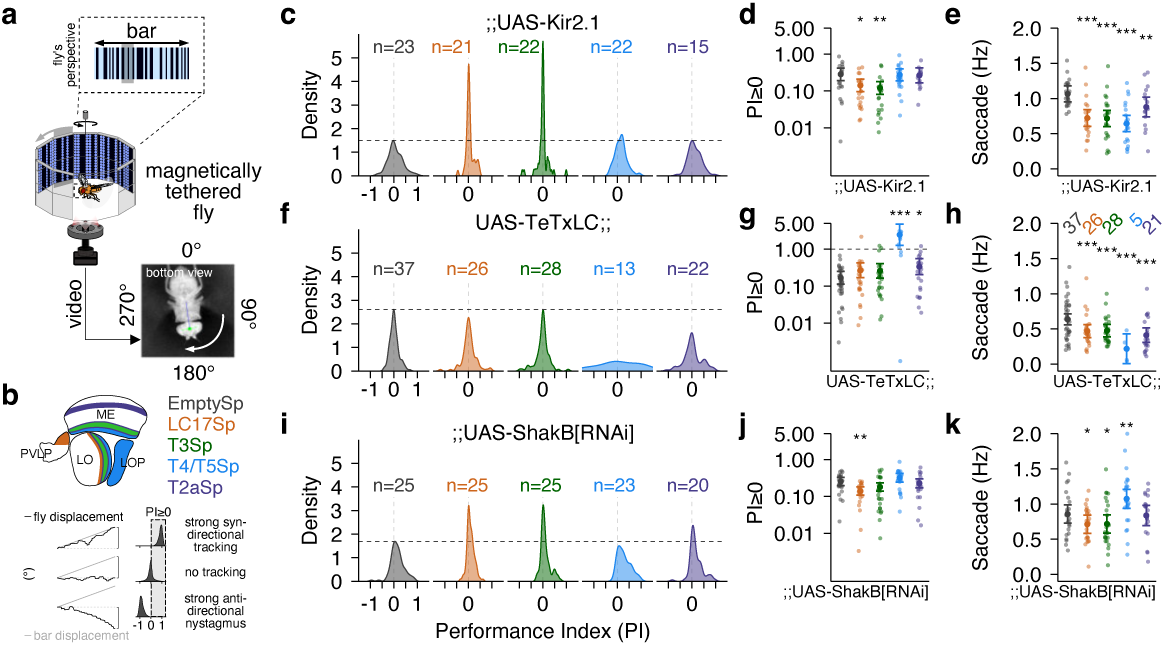
Gap junctional communication supports bar tracking in LC17 and T3 neurons. **a**, Top: representation of a motion-defined bar eliciting orienting behavior. Bottom: schematic illustrating a magnetically tethered fly free to rotate in the yaw plane within a surrounding LED display. Video recordings of the fly behavior were then tracked and its heading extracted for analysis. **b**, Top: schematic of the neuropils innervated by the driver lines tested; EmptySp (gray), LC17Sp (orange), T3Sp (green), T4/T5Sp (blue), and T2aSp (purple). Bottom: graphical representation of the tracking performance index (PI). **c**, PI distribution (kernel density estimation) in flies expressing Kir2.1 (EmptySp, n = 23, 4 trials; LC17Sp, n = 21, 4 trials; T3Sp, n = 22, 4 trials; T4/T5Sp, n = 22, 2 trials; T2aSp, n = 15, 4 trials). LC17 and T3 flies show a high peak around zero, meaning that they do not pursue the bar. Dashed horizontal black line indicates distribution peak in EmptySp flies. Data on EmptySp, T3Sp, and T4/T5Sp are replotted from Frighetto & Frye, 2023. **d**, Average PI values in flies expressing Kir2.1 that show PI greater or equal to zero, i.e., overall positive syn-directional tracking. Small opaque dots represent single flies. Bigger solid dots represent the sample mean and error bars ± 95% C.I. Data are shown in log scale. Statistical analysis was performed using unpaired two-sided t-test (*P<0.05, **P<0.01, ***P<0.001). **e**, Average frequency of body saccades in flies expressing Kir2.1. Small opaque dots represent single flies. Bigger solid dots represent the sample mean and error bars ± 95% C.I. Statistical analysis was performed using unpaired two-sided t-test (*P<0.05, **P<0.01, ***P<0.001). **f**, Kernel density estimation of PI distribution, as in **c**, in flies expressing TeTxLC (EmptySp, n = 37, 4 trials; LC17Sp, n = 26, 4 trials; T3Sp, n = 28, 4 trials; T4/T5Sp, n = 13, 4 trials; T2aSp, n = 22, 4 trials). LC17 and T3 flies pursue the bar as well as controls. **g**, Average PI values, as in **d**, in flies expressing TeTxLC that show PI greater or equal to zero. **h**, Average frequency of body saccades, as in **e**, in flies expressing TeTxLC. **i**, Kernel density estimation of PI distribution, as in **c**, in flies expressing ShakB[RNAi] (EmptySp, n = 25, 4 trials; LC17Sp, n = 25, 4 trials; T3Sp, n = 25, 4 trials; T4/T5Sp, n = 23, 4 trials; T2aSp, n = 20, 4 trials). LC17 and T3 flies show a relatively high peak around zero, meaning that they do not pursue the bar very well. **j**, Average PI values, as in **d**, in flies expressing ShakB[RNAi] that show PI greater or equal to zero. **k**, Average frequency of body saccades, as in **e**, in flies expressing ShakB[RNAi]. Driver lines are color-coded as in **b**.

To investigate the type of neurotransmission involved in this visual circuit, we blocked synaptic release of neurotransmitters in T3 and LC17 using the tetanus toxin light chain (TeTxLC^70^). Surprisingly, this manipulation in T3 and LC17 had no effect on either bar tracking performance (Fig. 3f,g) or optomotor gain (Extended Data Fig. 4c), although saccade frequency was reduced (Fig. 3h). Positive controls verified this manipulation: TeTxLC expression in T4/T5 abolished optic-flow stabilization (Extended Data Fig. 4c). These results indicate that hyperpolarization of LC17 impaired bar tracking behavior, but chemical synaptic blockade did not. This observation led us to query the role of electrical transmission.

Wide-field selective VPNs of the lobula plate have long been known to sculpt complex RFs via dendro-dendritic gap junction signaling^71–74^. We therefore considered whether the divergent Kir2.1 and TeTxLC phenotypes could be explained by electrical synapses (gap junctions^75^) that would respond to hyperpolarization but not chemical synaptic blockade. A recent study mapped the distribution of all innexin types in the fly brain at the light level^74^, identifying Shaking B (*shakB*, also known as innexin 8) to be the only gap junction protein expressed in neurons with broad distribution across all brain neuropils^74,76^. Hence, we hypothesized that T3 and LC17 neurons communicate via gap junctions. To test this, we used the bar tracking behavioral assay with RNA interference (RNAi^77^) to knockdown *shakB*^74^. We validated the efficacy of RNAi reagent using single-molecule HCR-FISH (Hairpin Chain Reaction Fluorescent In Situ Hybridization^78^); the number of *shakB* transcripts was significantly reduced in the cytoplasm of LC17 neurons (Extended Data Fig. 5a,b). Pan-neuronal *shakB* knockdown produced severely abnormal flight and leg posture as previously characterized in *shakB* mutants^79^ (Extended Data Fig. 4e). Knockdown of *shakB* selectively in T3 and LC17 neurons reduced bar tracking performance (Fig. 3i), with particularly severe effects in LC17 neurons, where even flies that managed to pursue the bar at all showed weak performance (Fig. 3j). Saccade frequency was also mildly decreased (Fig. 3k). These results indicate that electrical transmission is sufficient for bar tracking behavior and complement the TeTxLC results, where synaptic blockade reduced saccades but not tracking performance (this result requires a strong-acting variant of TeTxLC, see Extended Data Fig. 4g). Again, these effects are specific to object approach behavior; *shakB* knockdown in either LC17 or T3 preserved optomotor gain and avoidance responses, whereas its removal from T4/T5 reduced optomotor gain without impairing bar tracking (Extended Data Fig. 4d,f). In T2a neurons, *shakB* knockdown specifically reduced avoidance of small revolving squares and the frequency of associated saccades (Extended Data Fig. 4f). Together, these findings suggest that gap junctions in T3 and LC17 are crucial for peripheral bar discrimination computations required for sustained tracking, whereas chemical synapses seem to play a peripheral role in saccade initiation.

### *shakB* expression defines dendritic coupling in LC17 neurons

To assess *shakB* expression in LC17 and T3 neurons, we used the *shakB*-Trojan-Gal4 gene trap line^74,80^, which drives Gal4 under the control of the endogenous *shakB* regulatory elements (Fig. 4a, left). This strategy effectively reports the transcriptional state of the *shakB* locus, visualizing cells with active *shakB* expression. We examined colocalization between this reporter and cell-type-specific driver lines (LC17- and T3/T2a-LexA). In LC17, the overlap was prominent within the LC17 glomerulus as well as in lobula layer 2 and 3 (Lo2-3) where LC17 dendrites arborize (Fig. 4a, middle). By contrast, interpreting overlap in T3 was more challenging because the driver we used labels both T3 and T2a neurons. We observed a cluster of *shakB*-positive cell bodies that are intermingled in the region posterior-lateral to the lobula plate, where the somata of T3, T2a, and T2 reside (Fig. 4a, left). Consistent with this, we observed reporter overlap with T3/T2a-LexA in Lo2-3 layers, where both project axons, and in medulla layer 9 (M9), where both arborize dendrites (Fig. 4a, right). Single-cell RNA sequencing (scRNA-seq) datasets provided clearer evidence for cell-type-specific *shakB* expression. Using a developmental atlas of the optic lobe^25,81^, we identified a cluster containing LC12 and LC17 neurons (based on the expression of known marker genes^82^) with a very high expression of *shakB* relative to most other optic lobe neuronal types (Fig. 4b and Extended Data Fig. 6a).

**Figure 4.**
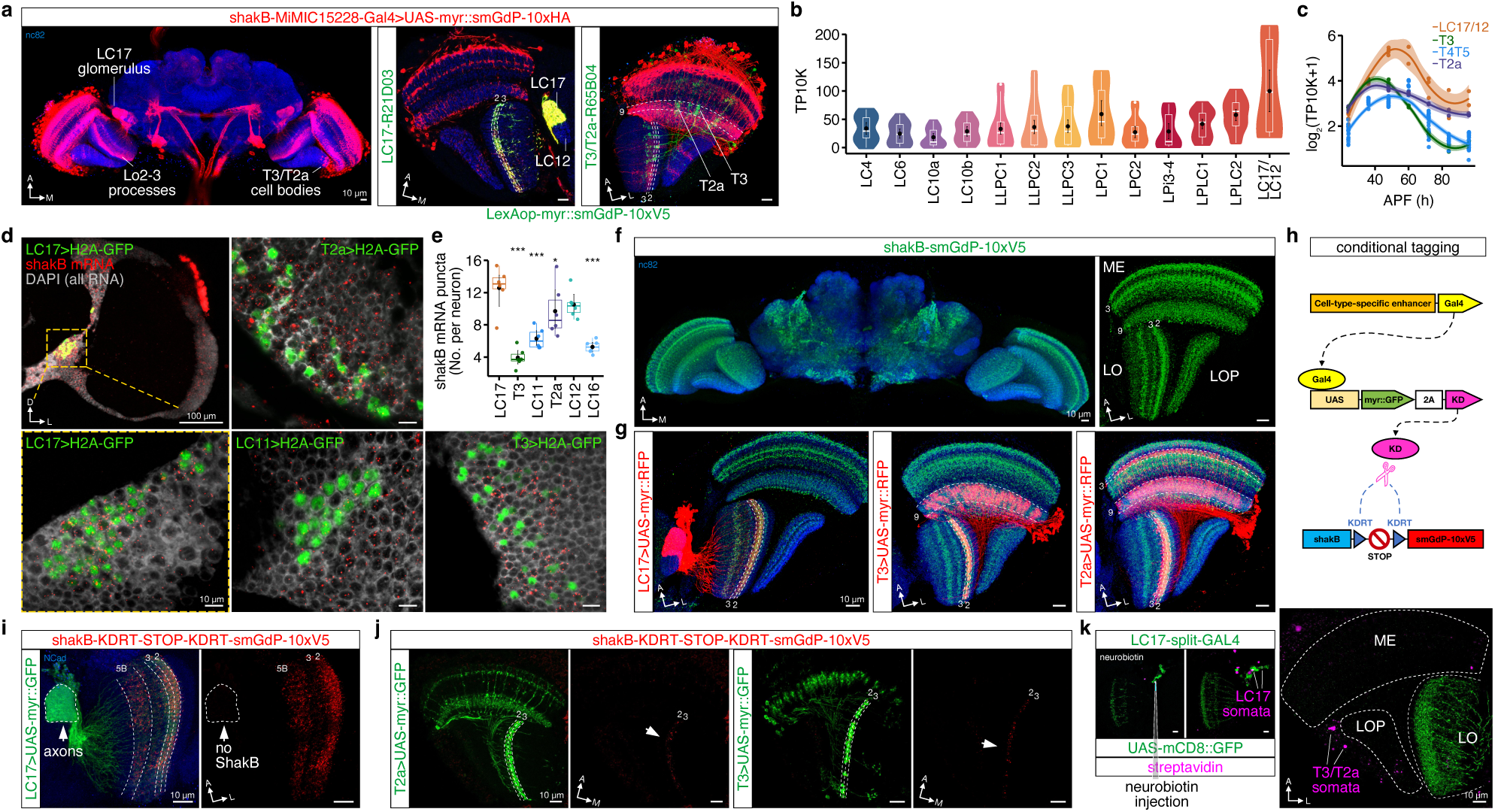
*shakB* expression and localization in LC17 and T3 neurons. **a**, Left: confocal projection of *shakB*-Trojan-Gal4 driver line (Ammer et al., 2022), highlighting LC17 glomerulus, innervations of Lo2-3, and cell bodies of T3, T2a, and T2. Middle: co-localization of *shakB*-Trojan-Gal4 and LC17 neurons. Overlap is highlighted in the LC17 glomerulus and Lo2-3. Right: co-localization of *shakB*-Trojan-Gal4 and T3/T2a neurons. Scale bar, 10 µm. **b**, Violin-box plots of the normalized expression level (TP10K, Transcript Per 10,000) in neurons innervating lobula and/or lobula plate across development (Kurmangaliyev et al., 2020). Black dots and error bars represent mean and bootstrapped ±95% C.I. **c**, Developmental expression pattern of *shakB* in LC17/LC12 (orange), T3 (green), T4/T5 (blue), and T2a (purple) (Kurmangaliyev et al., 2020), log-scaled normalized expression level. Dots represent replicates, lines represent smoothed conditional means with loess (local regression) method, and error bands ± 95% C.I. **d**, Top-left: light-sheet projection of the optic lobe showing LC17 nuclei and transcripts of *shakB*. Scale bar, 100 µm. Bottom-left: single 0.5 µm thick slice from panel on the top (dashed yellow box). LC17 cell bodies show high level of *shakB* transcripts. Scale bar, 10 µm. Top-right: same for T2a neurons. Bottom-middle: same for LC11 neurons. Bottom-left: same for T3 neurons. **e**, Box plots of *shakB* puncta counts in neurons from **d**. Colored dots represent averaged expression across all cells in single flies (n = 6, one brain side per animal). Black dots and error bars represent mean and bootstrapped ±95% C.I. Unpaired two-sided t-test between LC17 and other neurons (*P<0.05, **P<0.01, ***P<0.001). **f**, Left: confocal projection of constitutively tagged *shakB* allele. Right: expression pattern in the optic lobe. Scale bar, 10 µm. **g**, Left: confocal projection of co-localized constitutively tagged *shakB* allele and LC17 neurons marked with a membrane-bound reporter. Overlap is highlighted in Lo2-3. Middle: same for T3 neurons. Right: same for T2a neurons. **h**, Schematic of the conditional tagging approach. **i**, Confocal projection of conditionally tagged *shakB* allele in LC17 neurons. Scale bar, 10 µm. **j**, Left: same as in **i** for T2a neurons. Right: same as in **i** for T3 neurons. Scale bar, 10 µm. **k**, Left: injection of neurobiotin during whole-cell patch-clamp recording in a LC17 neuron (white). Gap junction-impermeable dye Alexa 594 was included to identify the filled neuron (white). Neighboring LC17 and other unidentified neurons are labeled after staining against neurobiotin with streptavidin (magenta). Right: maximum intensity projection of somata from putative T3, T2a, or T2 neurons (magenta) showing dye-coupling with the LC17 (not visible in these z-planes). Scale bar, 10 µm.

LC17/LC12, T3, T2a, and T4/T5 neurons all exhibited a pronounced peak of *shakB* expression around 48 h after puparium formation (APF, Fig. 4c), but in adult flies (i.e., 96 h APF), *shakB* expression declined sharply in T3 and T4/T5 neurons. By contrast, T2a retained modest levels of *shakB*, while LC17/LC12 sustained strong expression into adulthood (Fig. 4c). To validate the scRNA-seq data while preserving anatomical context, we used HCR-smFISH combined with Expansion-Assisted Light Sheet Microscopy (ExLSM^83^) to visualize and count individual *shakB* transcripts within identified somata (Fig. 4d). In adults, LC17, LC12 and T2a neurons displayed markedly higher *shakB* expression compared to T3 and to negative controls such as LC16 and LC11 (Fig. 4d,e and Extended Data Fig. 6b). However, gene expression data alone is insufficient to demonstrate gap junction coupling, since distinct neurons could restrict ShakB protein to different compartments; for example, T2a or T3 neurons might be receiving electrical inputs from upstream partners in the medulla, and LC17 neurons might instead connect electrically to downstream targets in the PVLP. To resolve this, we extended the RNA-level findings to protein and examine the subcellular localization of ShakB. We introduced an smGdP-10xV5 epitope tag into the endogenous *shakB* locus, enabling antibody-based detection of ShakB while preserving its native localization and expression levels^84^. We generated both constitutive (whole-animal) and conditional (cell-type-specific) versions of this tagged allele. In the constitutive allele (Fig. 4f), ShakB was most abundant in Lo2-3 layers, overlapping with the targeting region of LC17 dendrites and the axons of T2a and T3 neurons (Fig. 4g and Extended Data Fig. 6c). Using the conditional allele, we mapped subcellular localization of ShakB in specific cell types (Fig. 4h). LC17 neurons displayed ShakB puncta in their dendrites but not in their axons (Fig. 4i), indicating that gap junctional inputs converge on the dendrites. As a control, LC4 neurons showed puncta in both dendrites (Lo1-2 and Lo4 layers) and axons (Extended Data Fig. 6d). In T2a neurons, the puncta localized to Lo2-3 layers where they send presynaptic outputs; likewise, in T3 neurons the puncta were restricted to Lo2-3 (Fig. 4j). To note, in T4/T5 neurons ShakB puncta were restricted to the postsynaptic layers (i.e., M10 and Lo1) supporting the optomotor effect in the knockdown experiments (Extended Data Fig. 6g). These findings establish that LC17 neurons localize ShakB to their dendrites, consistent with the formation of putative gap junctions with coupling partners that may include other LC17 and upstream neurons such as T3 and T2a. To demonstrate functional gap junction coupling, we injected neurobiotin, a gap-junctions-permeable tracer, into patched-clamped LC17 neurons (Fig. 4k). The dye spread into other LC17 neurons as well as into cells with somata adjacent to the posterior-lateral lobula plate (i.e., matching the T3, T2a, and T2 somata location) (Fig. 4k and Extended Data Fig. 6e).

### Interplay between chemical and electrical synapses

Electrical coupling between T3 and LC17 neurons inferred from our molecular analysis corroborates the behavioral data (Fig. 3i-k). However, *shakB* knockdown in T3 and LC17 impaired overall bar tracking performance, but only marginally affected the frequency of body saccades. At the same time, synaptic blockade strongly reduced the frequency of body saccades without affecting the overall tracking performance. This suggests that the number of saccades *per se* is not a predictor of tracking performance and that their dynamic variables such as amplitude might also contribute. The relatively low saccade rate produced by the knockdown of gap junctions could have been combined with perturbed saccade dynamics leading to insufficient performance. By contrast, the low saccade rate resulting from blocking the chemical synapses could have been compensated with saccades of larger amplitude leading to sufficient performance.

We posit that these complex interactions between electrical and chemical synaptic manipulations combine when the T3-LC17 circuit is hyperpolarized, which reduces bar tracking performance by way of reduced saccade rate (Fig. 5a). To guide hypotheses about how early object vision might be modulated by gap junctions, we proposed a parsimonious conceptual model of T3-LC17 mixed synaptic function^75^. A visual object composed of ON and OFF edges robustly activate T3 feature detectors^35,54^ in a retinotopic manner (Fig. 5b). Depolarization of T3 membrane potential results in neurotransmitter release that generates an excitatory postsynaptic potential (EPSP) in LC17, which in turn translates in a calcium transient (EPSCaT). The latter, upon reaching threshold, triggers a body saccade as previously postulated^31,35^. Gap junctions between T3 and LC17 neurons, and among LC17 copies, transmit a coupling potential (assuming homotypic channels^76^) composed of a fast depolarization (i.e., spikelet) and subsequent hyperpolarization (i.e., afterhyperpolarization, AHP). In addition, gap junctions modulate the overall excitability of coupled cells by lowering input resistance^85^. It is reasonable to imagine that the subthreshold spikelet (i.e., fast depolarization) spreads postsynaptically across the network of coupled cells, generating a lateral excitation in LC17 neurons. This mechanism is fundamental for saccade dynamics, but it is not evident in the slow integration time of an EPSCaT. Yet, the AHP could last long enough to affect the response profile of an EPSCaT. The result of combining chemical and electrical transmission is a sharpened EPSCaT in LC17 with both reduced onset amplitude (because of decreased input resistance) and enhanced offset decay (because of the AHP). In this scenario, downregulation of *shakB* in LC17 would: (1) increase both the amplitude and duration of EPSCaTs (because of the sole activity of the excitatory chemical synapses); (2) remove the tight spatiotemporal activation and produce perturbed saccade dynamics (because of perturbed EPSCaTs and the lack of lateral excitation) (Fig. 5b). Next, the neuronal computations made by T3 and LC17 would presumably be integrated with other visual circuitry and filtered by a biophysical cascade of inter- and descending neurons, muscular excitation-contraction dynamics, skeletal kinematics, and aerodynamics.

**Figure 5.**
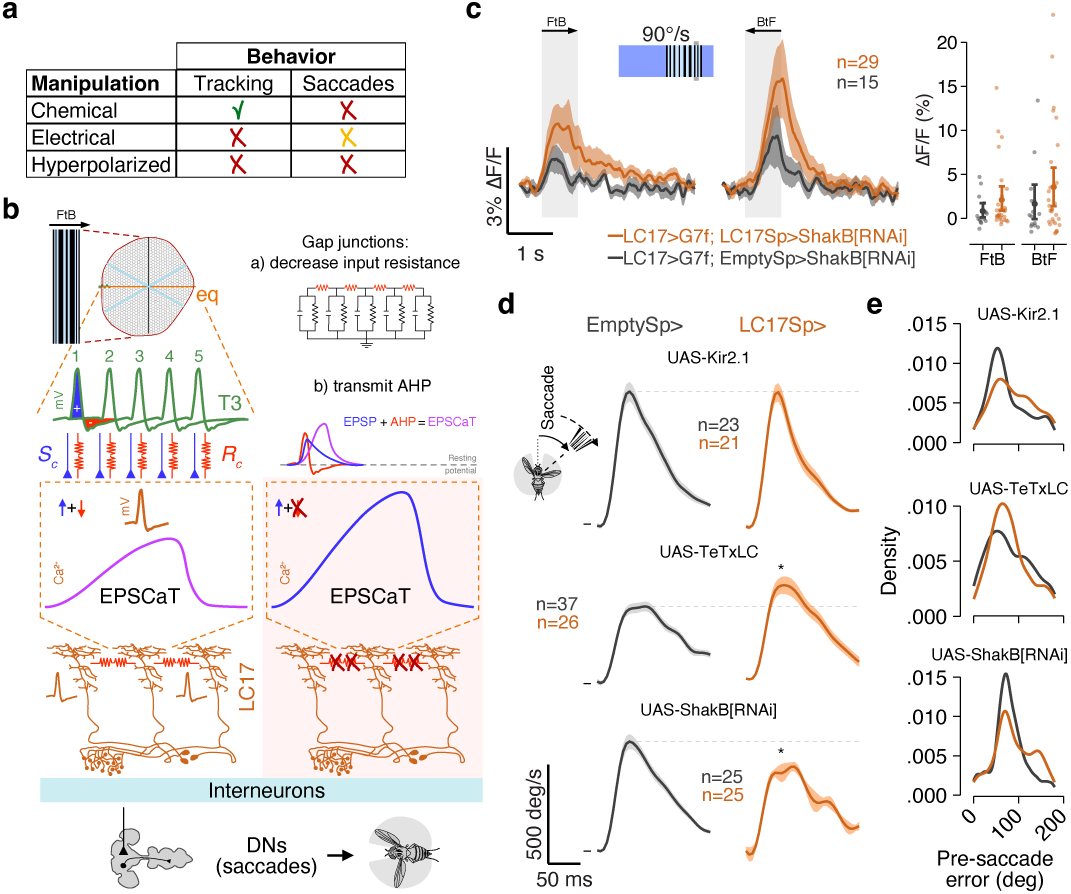
Gap junctions sculpt the signal for body saccade dynamics. **a**, Table summarizing the behavioral results relative to synaptic manipulations. **b**, Left: model of gap junctions’ role. A visual stimulus (motion-defined bar) moves FtB and starts to stimulate the photoreceptors housed inside the ommatidia. For simplicity, the fly eye is restricted to a single one-dimensional layer along the equator (eq, orange line). By skipping some intermediate connections within lamina and medulla, the information from the photoreceptors is then transmitted to T3 neurons (green). Postsynaptically, LC17 neurons accumulate the inputs resulting in an excitatory postsynaptic calcium transient (EPSCaT, purple curve). The EPSCaT is subject to two opposite mechanisms. In one, the excitatory chemical synapses (*S_c_*, blue triangles) increase the calcium accumulation. In the other, the gap junctions (*R_c_*, red resistance) decrease the membrane potential by lowering input resistance and by transmitting afterhyperpolarization (AHP). Consequently, this affects the EPSCaT by reducing onset amplitude and offset decay. Concurrently, gap junctions transmit fast depolarizations that excite and synchronize the circuit (i.e., lateral excitation). Right: model predictions of gap junctions down-regulation. A larger and longer EPSCaT would be generated. As a consequence, this disrupted calcium dynamics distorts the dendritic computation and the signal sent to the descending neurons that tune the shape of visually evoked body saccades. **c**, Right: average calcium traces in response to motion-defined bars moving FtB and BtF at 90°/s. ROIs drawn around LC17 dendrites in control (gray; n = 15) and ShakB[RNAi] (orange; n = 29) flies. Right: average peak response (200 ms window) in controls and ShakB[RNAi] to a motion-defined bar. Small opaque dots represent single average peak responses. Bigger solid dots represent the sample mean and error bars ± 95% C.I. **d**, Average angular velocity during directed and syn-directional body saccades (mean ± s.e.m.) in EmptySp (gray) and LC17Sp (orange) flies. From top to bottom: flies expressing Kir2.1, TeTxLC, and ShakB[RNAi]. Statistical analysis on peak angular velocity (25 ms after saccade initiation) was performed using unpaired two-sided t-test (*P<0.05, ***P<0.001). **e**, Kernel density estimation of the pre-saccade error angle in flies expressing Kir2.1 (top), ShakB[RNAi] (middle), and TeTxLC (bottom). Driver lines are color-coded as in **d**.

To explore these predictions, we performed calcium imaging and extracted the behavioral parameters from previously collected data in *shakB* RNAi expressing flies. First, the calcium responses to a motion-defined bar moving at a speed close to the one used in behavioral experiments were higher and wider in flies with *shakB* knockdown in LC17 compared to controls (Fig. 5c). Second, the peak angular velocity during saccades was reduced in *shakB* knockdown compared to controls (Fig. 5d). Notably, the peak angular velocity was increased in flies with synaptic blockade, confirming the compensatory mechanism for reduced saccade rate (Fig. 3h and 5d). Finally, the angular distance between fly heading and bar position before a saccade was triggered (i.e., pre-saccade error angle) showed larger angular distances in *shakB* RNAi expressing flies and more spatially restricted distances in TeTxLC expressing flies compared to controls (Fig. 5e). This means that the compromised saccade dynamics caused by the knockdown of gap junctions indirectly affects the timing for triggering saccades, and consequently their number, during bouts of tracking (Extended Data Fig. 8). In short, we posit that T3 and LC17 rely on mixed chemical and electrical synapses to enable proper behavioral dynamics during orientation; chemical synapses in LC17 neurons trigger saccades while gap junctions tune their dynamics. This simple conceptual model captures features underlying the behavior and physiology of visual approach.

## Discussion

We identified LC17 visual projection neurons as a key downstream partner of T3 neurons, forming a visual pathway that supports object tracking during flight. Using connectomics, physiology, behavior, anatomy, and molecular genetics, we showed that LC17 neurons receive excitatory input from T3, exhibit tuning properties well suited for bar tracking, and are necessary for orienting behavior. We demonstrated that electrical synapses mediate LC17’s role in tracking. Consistent with this, LC17 neurons express high levels of *shakB*, localize ShakB protein to their dendrites, and show functional electrical coupling both with other LC17 neurons and upstream partners (i.e., T3 and T2a neurons). These findings define a new pathway for the visual control of approach behavior, while also revealing a central role for gap junctions in guiding rapid visual orientation behavior.

The tuning properties of LC17 neurons position them to support selective object pursuit. Unlike LC11 (another postsynaptic partners of T3 neurons), which detect small objects and are suppressed by wide-field stimuli, LC17 respond to bars, integrate ON and OFF contrasts, and exhibit broad edge sensitivity^42,45,46^. Their rapid adaptation and decreased responses with repeated stimuli suggest a role in saliency detection, enabling the system to prioritize novel or dynamic features over common revisited ones. Direct connections between LC and descending neurons that regulate visuomotor transformation for escape responses^8^ exemplify the linear computations underlying avoidance^86^ (Extended Data Fig. 9a). For instance, two looming stimuli presented from opposite sides simultaneously must be averaged for the fly to escape from both. By contrast, orientation for approach requires nonlinear computations to select one object and filter out others^87^, a winner-take-all strategy^88,89^. Thus, the underlying neural circuits for approach may involve recurrent networks and less streamlined feedforward connections with premotor centers, which appears to be the case with LC17 (Extended Data Fig. 9b,c). Broad visual and optogenetic stimulations did not activate LC17, consistent with a mechanism of global competitive inhibition – as described in vertebrate optic tectum^88–90^ – that enforces selection of a single target. The approach circuit may combine a bottom-up component that detects relevant visual objects and suppresses unrelated stimuli through wide-field inhibitory interneurons (e.g., LT56-Glu and mALC2-GABAergic^21^), with a top-down component that modulates circuit sensitivity. Notably, centrifugal octopaminergic neurons such as AL2i2 ramify near Lo2-3 layers, where LC17 dendrites and T3/T2a axons arborize, as well as in M9 layer, where T3/T2a dendrites reside^91^. Given the facilitative effects of octopamine on the fly visual circuits^92–95^ and the high expression of octopamine receptors in LC17 (Yerbol Kurmangaliyev, unpublished data), this circuit motif appears plausible, although the net effect of neuromodulation remains difficult to predict^96^. Future studies will test these predictions and refine our understanding of the circuitry dependent upon LC17.

A central feature of this pathway is its reliance on electrical neurotransmission. Hyperpolarizing LC17 impaired tracking performance, but blocking chemical synaptic release did not, whereas *shakB* knockdown reduced bar pursuit. Together, these results show that gap junctions in LC17 are essential for sustained object tracking and regulate the timing of bar-directed body saccades. By contrast, chemical synapses from LC17 proved important for saccade initiation, as their blockade reduced the frequency of body saccades. These complementary roles suggest that mixed electrical-chemical transmission^75^ could endow the circuit with speed and flexibility; electrical coupling rapidly transmits retinal object position while compressing calcium transients trigger steering actions discretely. LC17 were recently shown to possess the highest number of mitochondria among LCs^97^, which might fulfill the high metabolic demand required by mixed electrical-chemical synapses for rapid communication as has been proposed for wing sensory and premotor neurons^98^.

Our analysis revealed robust *shakB* transcription and ShakB protein localization in LC17 dendrites, consistent with upstream electrical coupling. Although our dye filling experiment could not unequivocally identify specific types of T-cells, behavioral and protein localization data support electrical coupling between LC17 and the upstream partners T2a and T3 neurons. T2a neurons showed significant adult *shakB* expression and ShakB enrichment at presynaptic terminals, suggesting electrical coupling with LC17, but they did not seem important for bar tracking. This functional coupling might play a role in a different behavioral context or drive avoidance independently from LC17. On this note, electrical coupling with other neurons (e.g., LC12), represents an alternative hypothesis. Future investigations will further explore these issues.

Prior studies have shown that even a low level of ShakB is sufficient to sustain gap junctional communication in lobula plate tangential cells^99^ and in flight motoneurons^100^. A related scenario may apply to T3 neurons; although adult *shakB* expression was low, its knockdown disrupted bar tracking, indicating that relatively small amounts of ShakB are functionally important. *shakB* expression peaks broadly across many neuron types at ∼60 h APF^25^, then declines by eclosion (Extended Data Fig. 6g), indicating a universal development program (potentially related to spontaneous neural activity arising at the same time^101^). This may imply that early knockdown of *shakB* could elicit an effect on adult physiology and behavior by perturbing this developmental program and potentially altering the later maturation of chemical synapses^75,102^ or even affecting synaptic partner choice.

Synaptic blockade in LC17 did not impair bar tracking but reduced the frequency of body saccades, suggesting a division of labor: gap junctions at LC17 dendrites appear to be required to maintain sharp onset-offset dynamics, perhaps to localize T3-activated object positioning within the broad dendritic branches, while chemical synapses appeared to be required to evoke response amplitude sufficient to trigger saccades. A similar reduction in saccade frequency was also observed in T2a neurons, reinforcing their contribution to steering. Notably, this effect was sensitive to manipulation strength, as a weaker TeTxLC allele left both tracking performance and saccades intact (Extended Data Fig. 4g). Since the positional information needs to reach command neurons in the central brain, the absence of ShakB protein in LC17 axon terminals (i.e., glomerulus in the PVLP) suggests that dendritic electrical coupling with other VPNs (e.g., LT62 and LT66) may be functionally important. Alternatively, LC17 might be electrically coupled with downstream interneurons in the PVLP, but the protein in the axons was not detected by our conditional tagging. An important limitation of this approach is that it only reports protein levels after the driver lines start expressing the transcriptional activator (i.e., Gal4). In principle, this means that the homomeric hemichannels could already be localized in the junctional regions of presynaptic axons before the endogenous tagging begins to take effect. Indeed, by labelling the same neurons using driver lines that turn on at different timepoints the ShakB puncta can be visible in different neuronal compartments (Extended Data Fig. 7).

In conclusion, we identified a visual pathway for approach that relies on electrical coupling between T3 and LC17, and highlighted the functional role of *shakB* in establishing dendritic gap junctions in LC17. By integrating connectomics, genomics, genetics, physiology, and behavior, we reveal how electrical and chemical synapses contribute to rapid object detection and orientation. Future studies will determine how widespread such mixed transmission strategies are across the fly visual system and whether comparable mechanisms operate in vertebrate circuits for target selection and pursuit^103^.

**Extended Data Figure 1 (related to Figure 1).**
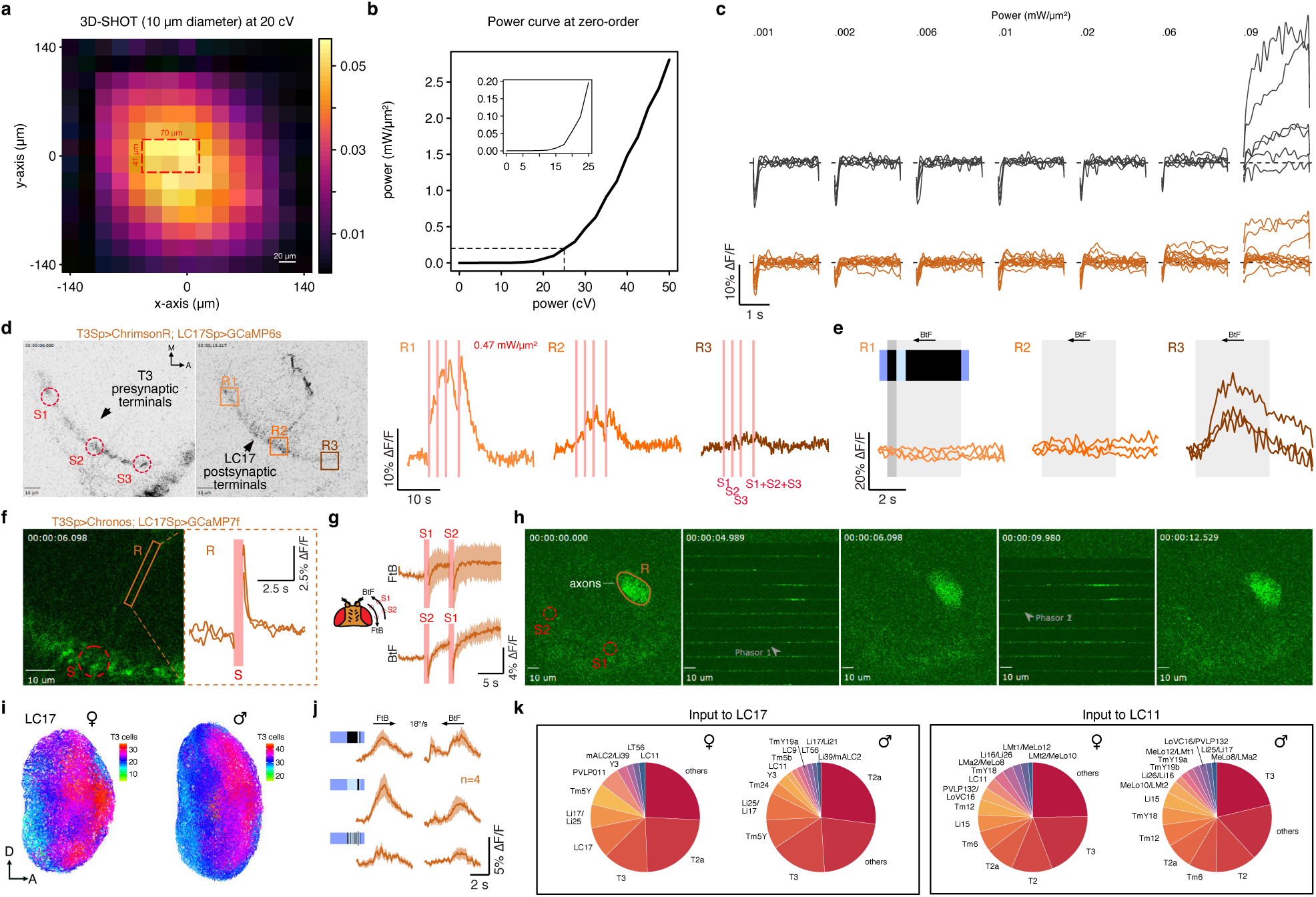
Complexity in functional response of LC17 neurons. **a**, 3D-SHOT power map computed by recording the operational range of the SLM with a camera placed below the objective. A single hologram was swept across the entire field of view (FOV) with steps of 20 µm and constant power. The light recorded from the camera was then calibrated by using a power meter. Red dashed rectangle represents the FOV used in our 3D-SHOT experiments. **b**, Power curve recorded using a power meter placed at the objective focal plane. Power increased by steps of 2.5 cV. Inset: close-up of the power curve below cell damage threshold. **c**, Calcium responses in LC17 dendrite to a single 3D hologram at increasing power in control flies (top row; gray; n = 6) and in flies expressing Chrimson in T3 (bottom; orange; n = 7). Stimulation duration, 0.5 s; Pulse frequency, 50 Hz (single pulse duration, 5 ms); ISI, 3 s. Above 0.06 mW/µm^2^ the calcium responses show sign of cell damage characterized by high increasing activity. **d**, Left: two-photon image of T3 presynaptic terminals expressing ChrimsonR tagged with tdTomato with three holograms (S1-3, red dashed circles) targeting them. Middle: ROIs drawn around LC17 postsynaptic terminals (R1-3, orange/brown squares) expressing GCaMP7f. Right: calcium traces in the three ROIs in response to a sequential stimulation (S1-3, red shaded regions starting from the left) with a fourth stimulation encompassing all three of them simultaneously (n = 1; 1 trial). Traces are interrupted during the stimulation period due to the shutting off of the PMTs to prevent damage. Pulse duration, 0.5 s; ISI, 1.5 s and 2.5 s; Power, 0.47 mW/µm^2^. Scale bars, 10 µm. **e**, Calcium traces in the three ROIs in response to an ON solid bar (9° x 72°, width x height) moving back-to-front (BtF) at 18°/s (n = 1; 3 trials). Light gray shaded region represents the overall stimulation duration, while the dark gray shaded region represents the stimulus dwell time (∼0.5 s) within the ROI receiving inputs by a few T3 terminals (∼9 µm) which is similarly to duration (0.5 s) and size (10 µm diameter) of the 3D holograms. **f**, Left: two-photon image of LC17 dendrites expressing GCaMP7f with illustrated hologram (S, red dashed circle) targeting T3 presynaptic terminals expressing Chronos and recorded ROI from a single neurite (R, orange solid rectangle). Right: calcium response to the targeted stimulation (n = 1; 2 trials). Pulse duration, 0.5 s; Power, 0.47 mW/µm^2^. **g**, Left: illustration of the direction simulated by two sequential 3D holograms (FtB, front-to-back; BtF, back-to-front). S1-S2 simulates FtB motion, while S2-S1 BtF motion. Right: average calcium traces from an ROI drawn around LC17 axons in response to FtB and BtF stimulations (n = 1; 3 trials; mean ± s.d.). Pulse duration, 1 s; ISI, 4 s; Power, 0.47 mW/µm^2^. A preferred directional integration seems to take place within the glomerulus. **h**, Two-photon images of LC17 neurons expressing GCaMP7f during the sequential stimulations. ROI recorded (R, orange ellipse) drawn around LC17 axons. 3D holograms (S1 and S2, red dashed circles) targeting T3 presynaptic terminals expressing Chronos. Arrows depict the hologram (Phasor) positions. **i**, LC17 dendrites in the lobula (lateral view) color-coded according to the number of T3 neurons each LC17 neuron receives input from. A posterior-anterior gradient is visible in both FAFB (left) and MAOL (right) connectomes. D, dorsal; A, anterior. **j**, Average calcium traces in response to three different visual stimuli (ON, OFF, and motion-defined bars) moving FtB and BtF at 18°/s in flies previously stimulated with 3D-SHOT (corresponding to Fig. 1i). **k**, Left: pie charts of the input neurons to LC17 neurons based on the number of synapses in FAFB (left) and MAOL (right). Right: same as on the right for LC11 neurons.

**Extended Data Figure 2 (related to Figure 1).**
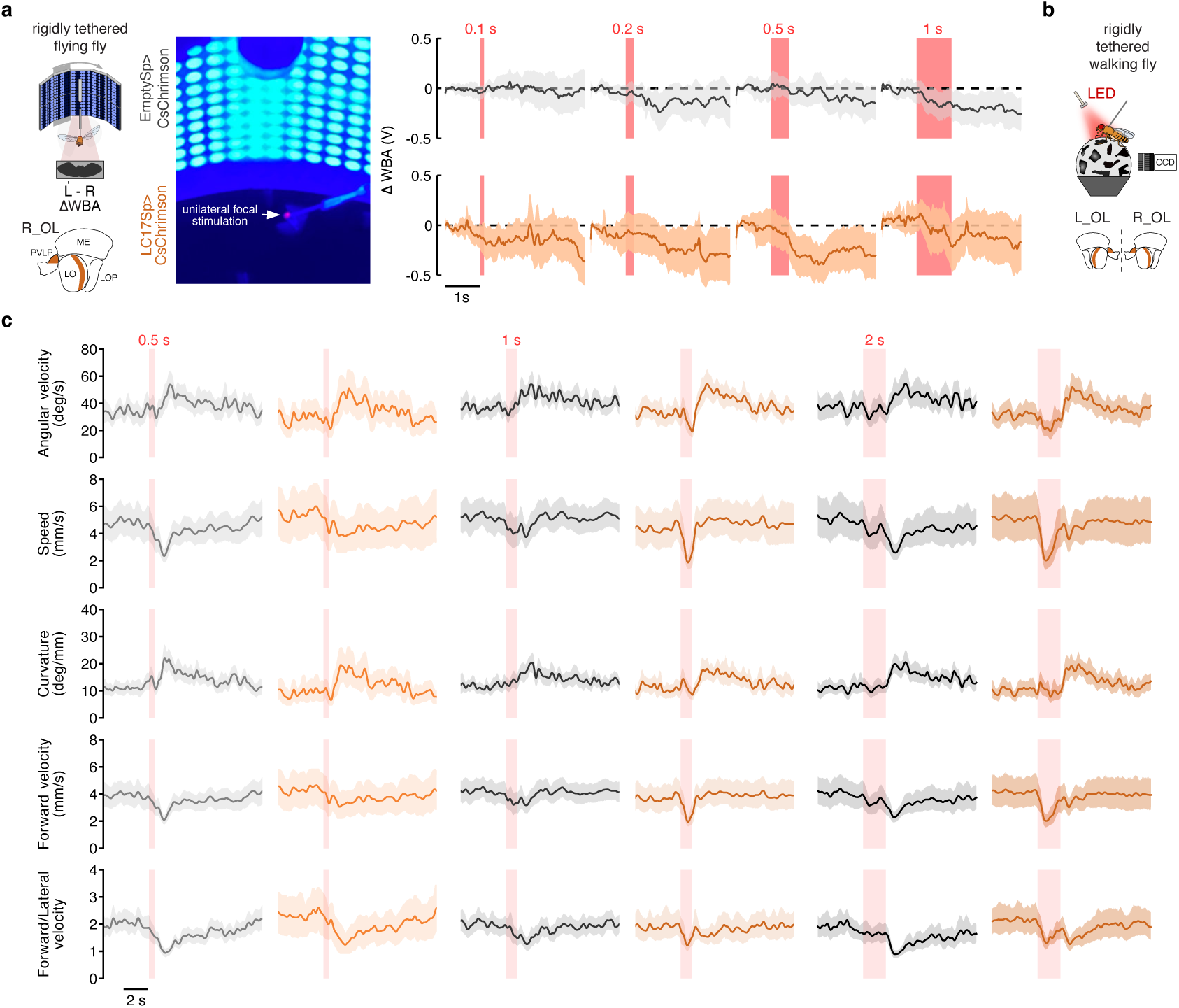
Behavioral response to LC17 ensemble optogenetic activation. **a**, Top-left: illustration of a rigidly tethered fly glued to a tungsten pin. An optical sensor placed underneath the fly records the difference between left and right wing beat amplitudes (ΔWBA) as a proxy of steering effort. Bottom-left: schematic of the region innervated by LC17 neurons (orange) in the right optic lobe (R_OL) and stimulated using optogenetic. ME, medulla; LO, lobula; LOP, lobula plate; PVLP, posterior ventrolateral protocerebrum. Middle: picture of the fly suspended within a surrounding LED display during unilateral focal stimulation over R_OL. Right: ΔWBA traces (mean ± s.e.m.) in EmptySp (n = 16; 1 trial) and LC17Sp (n = 15; 1 trial) flies expressing Chrimson in response to different stimulation durations (red shaded regions). **b**, Illustration of a rigidly tethered fly walking on an air-supported ball. A red LED for optogenetic was placed above the fly. **c**, Traces related to five kinematic variables (mean ± 95% C.I.) in EmptySp (n = 25-27; 10 trials) and LC17Sp (n = 13-25; 10 trials) flies expressing CsChrimson in response to different stimulation durations (red shaded regions) during walking.

**Extended Data Figure 3 (related to Figure 2).**
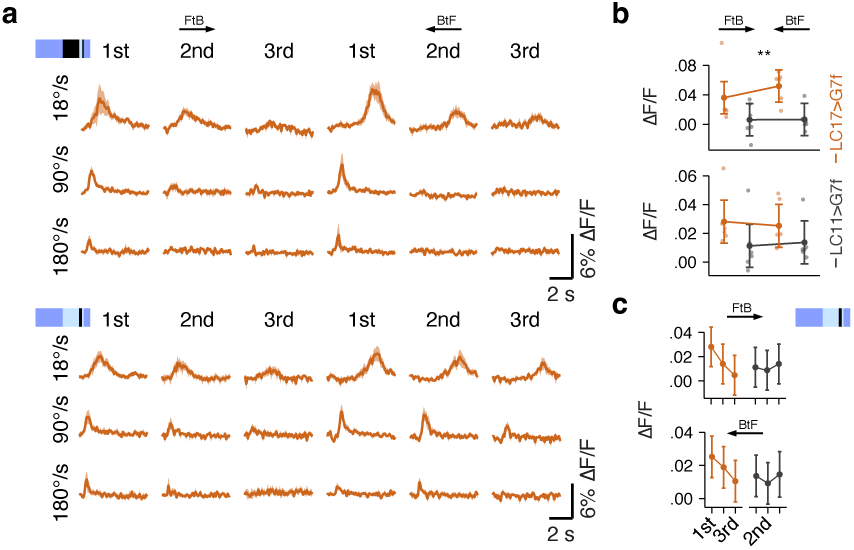
ON-OFF and directional sensitivity in LC17. **a**, Top: average calcium traces in response to an ON bar moving front-to-back (FtB) and back-to-front (BtF) at different velocities over three repetitions in LC17 (n = 6, mean ± s.e.m.). Bottom: same as on the top for a moving OFF bar. **b**, Top: maximum peak responses in LC17 (orange) and LC11 (gray) for FtB and BtF moving ON bar (9° x 72°, width x height) at 18°/s (mean ± 95% C.I.). Bottom: same as on the top for OFF bar. Statistical analyses were performed using unpaired two-sided t-test (*P<0.05, **P<0.01). **c**, Maximum peak response in LC17 (orange) and LC11 (gray) for FtB (top) and BtF (bottom) moving OFF bar at 18°/s over three repetitions.

**Extended Data Figure 4 (related to Figure 3).**
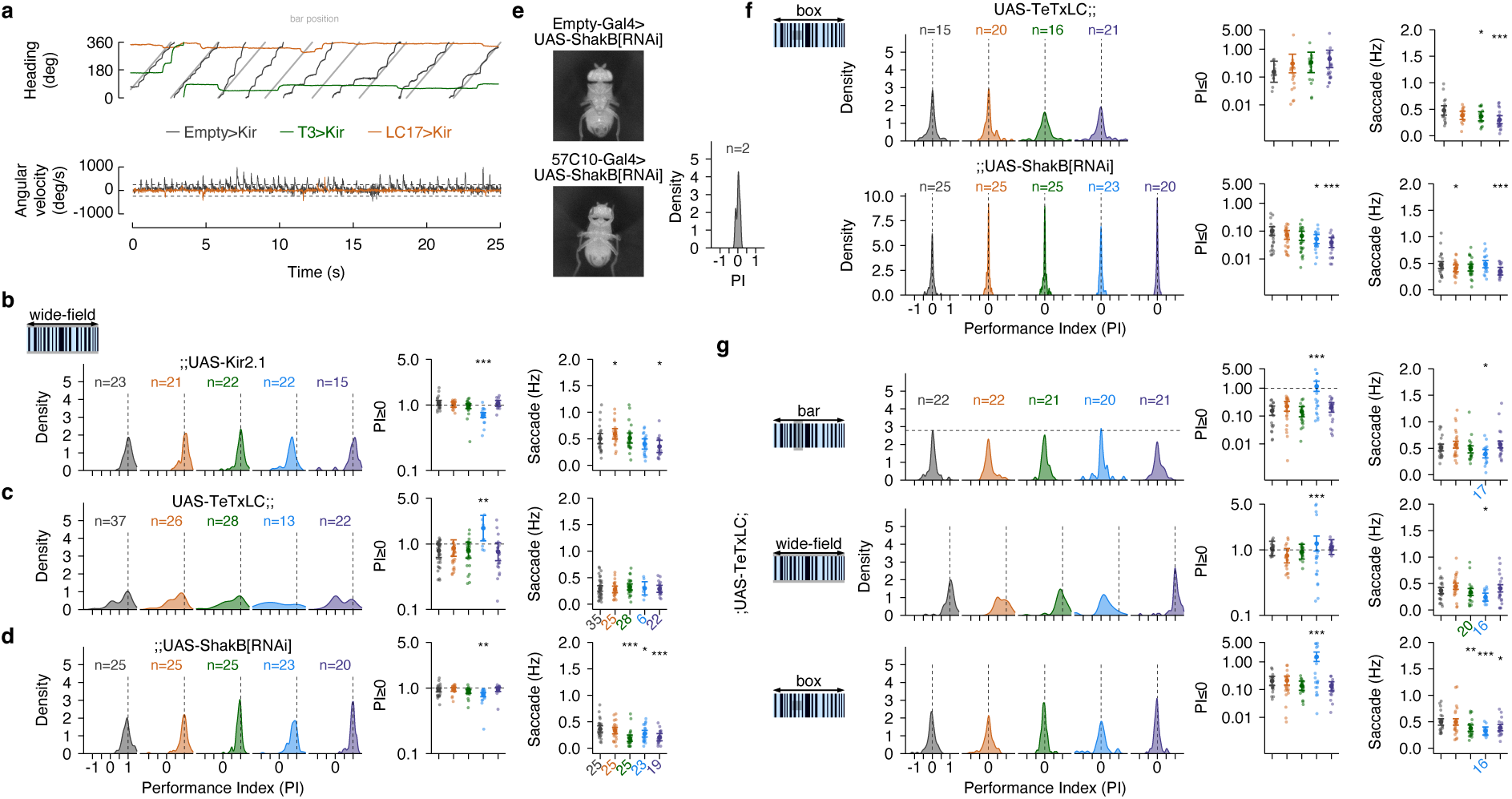
Manipulation of LC17 and T3 neurons does not affect the response to wide-field or small-objects. **a**, Top: wrapped heading traces during bar revolutions (light gray) at 112.5°/s for 25 s in LC17Sp (orange), T3Sp (green), EmptySp (gray) silenced flies. Bar position is representative and related to EmptySp. Bottom: angular velocity related to the traces on the top. Dashed horizontal black lines indicate the threshold for saccade detection. Data on EmptySp and T3Sp are replotted from Frighetto & Frye, 2023. **b**, Right: PI distribution (kernel density estimation of optomotor gain) in flies expressing Kir2.1 (EmptySp, n = 23, 4 trials; LC17Sp, n = 21, 4 trials; T3Sp, n = 22, 4 trials; T4/T5Sp, n = 22, 2 trials; T2aSp, n = 15, 4 trials). LC17 and T3 flies do not show a compromise response. Dashed vertical black line indicates a perfect optomotor gain (PI = 1). Data on EmptySp, T3Sp, and T4/T5Sp are replotted from Frighetto & Frye, 2023. Middle: average PI values in flies expressing Kir2.1 that show PI greater or equal to zero, i.e., overall positive syn-directional tracking. Small opaque dots represent single flies. Bigger solid dots represent the sample mean and error bars ± 95% C.I. Data are shown in log scale. Right: average frequency of body saccades in flies expressing Kir2.1. Small opaque dots represent single flies. Bigger solid dots represent the sample mean and error bars ± 95% C.I. **c**, Same as in **b** for TeTxLC expressing flies. **d**, Same as in **b** for ShakB[RNAi] expressing flies. **e**, Two exemplary flies with shakB knockdown in control (top) and pan-neuronal (bottom) flies. The fly on the bottom shows an abnormal leg position during flight typical of *shakB*[2] mutant flies (Homyk et al., 1980). As highlighted on the right, the bar tracking PI distribution is centered to 0, meaning that flies have a strongly compromised behavior. **f**, Top: same as in **b** for small square avoidance in TeTxLC expressing flies. Bottom: same as in **b** for small square avoidance in ShakB[RNAi] expressing flies. **g**, Same as in **b** for bar tracking (top), wide-field optomotor gain (middle), and small square avoidance (bottom) in a weaker version TeTxLC expressing flies. Statistical analysis was performed using unpaired two-sided t-test (*P≤0.05, **P<0.01, ***P<0.001).

**Extended Data Figure 5 (related to Figure 3).**
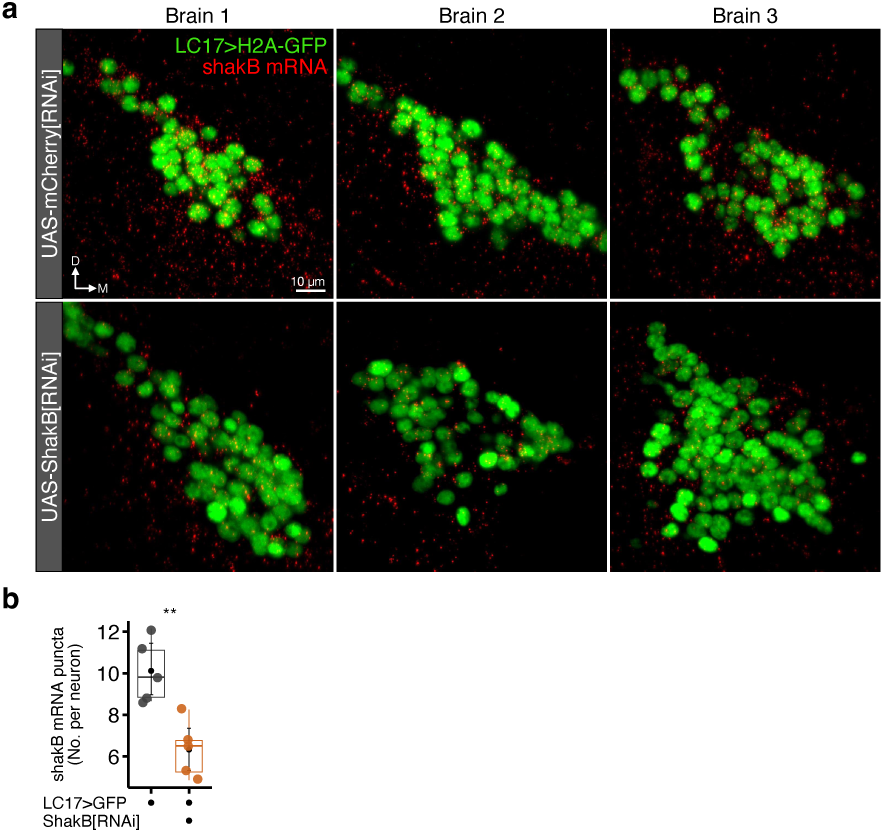
Validation of the UAS-ShakB[RNAi] effector line. **a**, Top: light-sheet single slice (0.5 µm) of LC17 nuclei (green) and transcripts of shakB (red) in control flies. Three exemplary brains are shown. Bottom: same as on the top for ShakB[RNAi] expressing flies. ShakB[RNAi] flies show low level of shakB transcripts around the nuclei compared to controls. Scale bars, 10 µm. **b**, Box plots of shakB puncta count in control (gray) and ShakB[RNAi] (orange) flies. Dilation, 50%; Brightness threshold, 10. Opaque dots represent average in single flies (n = 5). Black dots and error bars represent mean and bootstrapped ±95% C.I. Unpaired two-sided t-test between control and ShakB[RNAi] flies (**P<0.01).

**Extended Data Figure 6 (related to Figure 4).**
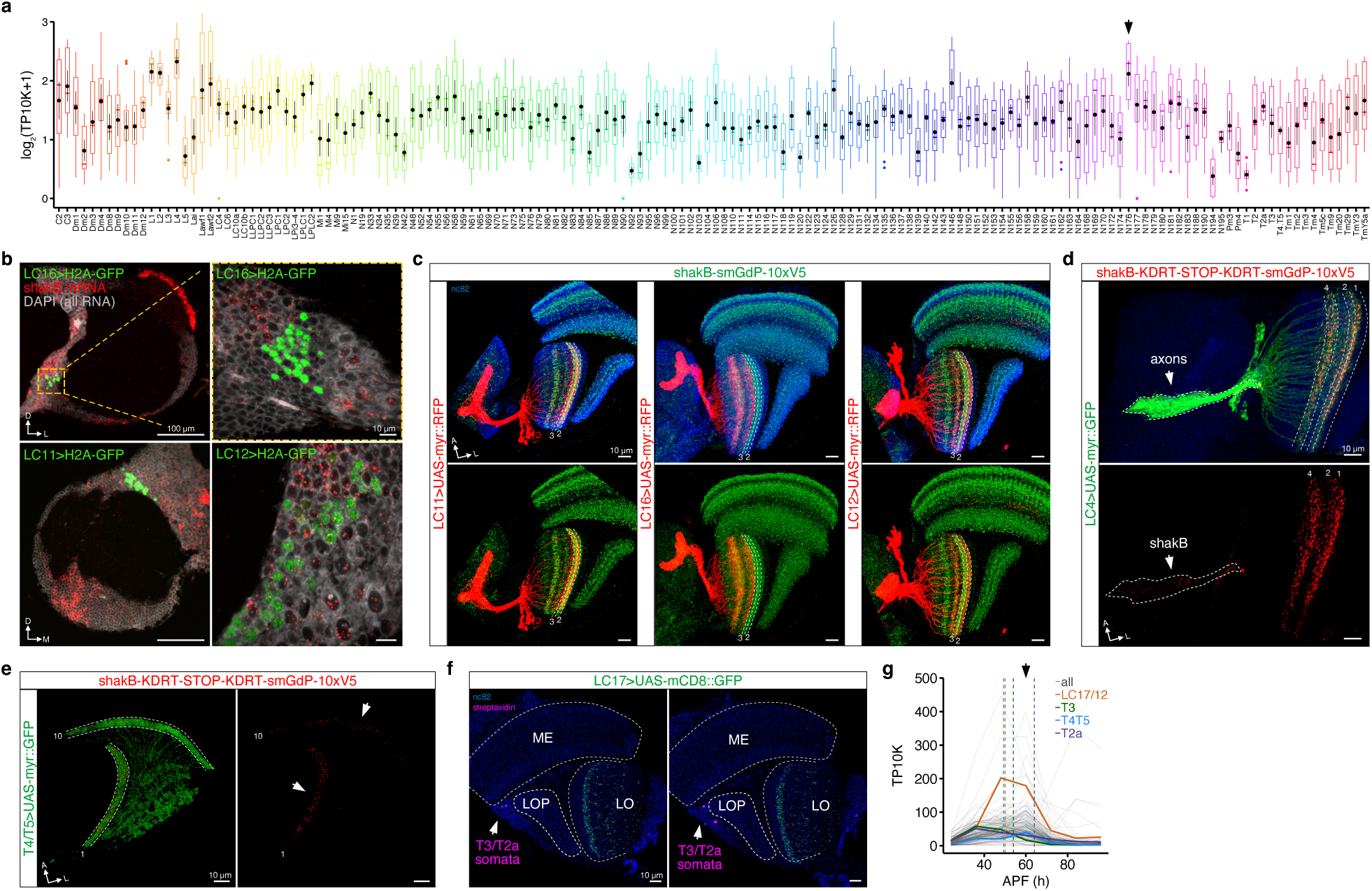
*shakB* expression in the optic lobe. **a**, Expression level of *shakB* in visual neurons across development (Kurmangaliyev et al., 2020). Data are shown in log-scaled normalized expression level (TP10K, Transcript Per 10,000). Black dots and error bars represent mean and bootstrapped ±95% C.I. N176 cluster, which includes LC17 and LC12, shows among the highest expression level (black arrow). **b**, Top-left: light-sheet projection of the optic lobe showing LC16 nuclei and transcripts of *shakB*. Scale bar, 100 µm. Top-right: single 0.5 µm thick slice from panel on the left (dashed yellow box). LC16 cell bodies show very low level of *shakB* transcripts. Scale bar, 10 µm. Bottom-left: optic lobe showing LC11 nuclei and transcripts of *shakB*. Bottom-right: slice showing a high amount of *shakB* transcripts in LC12 cell bodies. D, dorsal; L, lateral; M, medial. **c**, Left: confocal projection of co-localized constitutive *shakB* tagged allele and LC11 neurons. Some overlap is highlighted in Lo2-3. Middle: co-localization of constitutively tagged *shakB* allele and LC16 neurons. No overlap is identifiable. Right: same co-localization as before but with LC12 neurons. Weak overlap is visible in Lo2. **d**, Confocal projection of conditionally tagged *shakB* allele in LC4 neurons. ShakB is visible in dendrites (Lo1-2, and Lo4) and in axons. Scale bar, 10 µm. **e**, Same as in **d** but for T4/T5 neurons. ShakB is visible in postsynaptic terminals (M10 and Lo1). Scale bar, 10 µm. **f**, Somata of T3, T2a, or T2 neurons (magenta) show dye-coupling with the LC17. Scale bar, 10 µm. **g**, Average developmental expression of *shakB* in LC17/LC12 (orange), T3 (green), T4/T5 (blue), and T2a (purple), compared to all the other visual neurons (gray) from 24 to 96 h APF (Kurmangaliyev et al., 2020). Dashed vertical lines indicate the onset time for the corresponding split-Gal4 driver lines used (LC17, orange; T3Sp, green; T2a, purple). Black arrow indicates the developmental peak shows by the majority of visual neurons.

**Extended Data Figure 7 (related to Figure 4).**
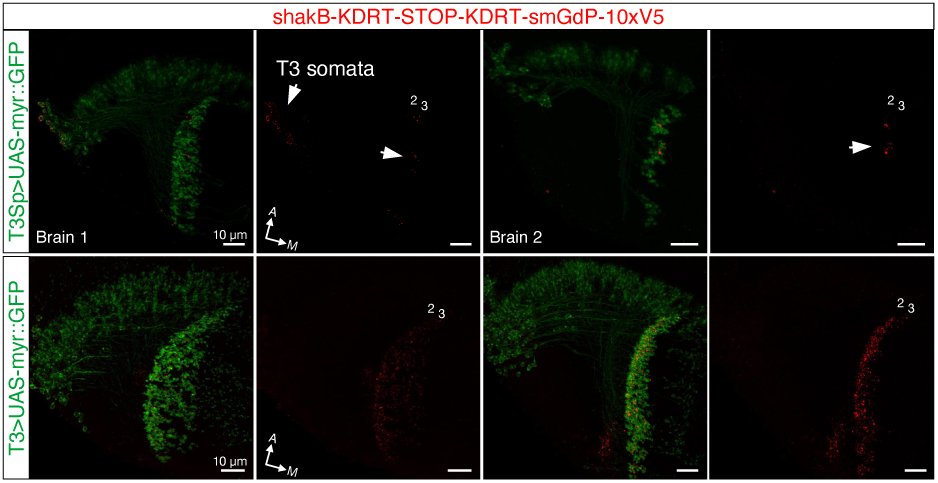
ShakB protein localization in T3 neurons. Top: confocal maximum intensity projection of conditionally tagged *shakB* allele in T3 neurons labelled by a split-Gal4 line (onset time, 54 h APF) in two brains. ShakB is visible in somata and/or very few presynaptic terminals (Lo2-3). Bottom: same as on the top for a Gal4 line (onset time, 48 h APF) which corresponds to the hemidriver (activation domain) of the split-Gal4. A Gal4 line with earlier onset shows ShakB accumulation within the presynaptic terminals (Lo2-3). Scale bars, 10 µm.

**Extended Data Figure 8 (related to Figure 5).**
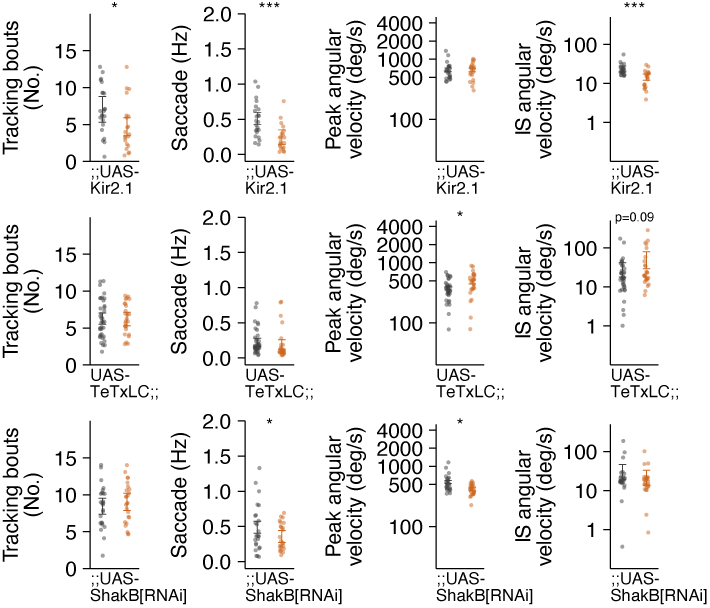
Parameters during bouts of tracking in EmptySp (gray) and LC17Sp (orange) flies. From left to right: Average of bouts of bar tracking per fly. Small opaque dots represent single flies and error bars ± 95% C.I. The bouts are defined as periods of tracking in which the R-squared captured by a linear regression is equal or greater than 0.75; Average frequency of body saccades; Average peak of angular velocity during saccades; Average inter-saccade velocity. Different rows represent flies expressing Kir2.1 (top), ShakB[RNAi] (middle), and TeTxLC (bottom). Statistical analyses were performed using unpaired two-sided t-test (*P<0.05, **P<0.01, ***P<0.001).

**Extended Data Figure 9.**
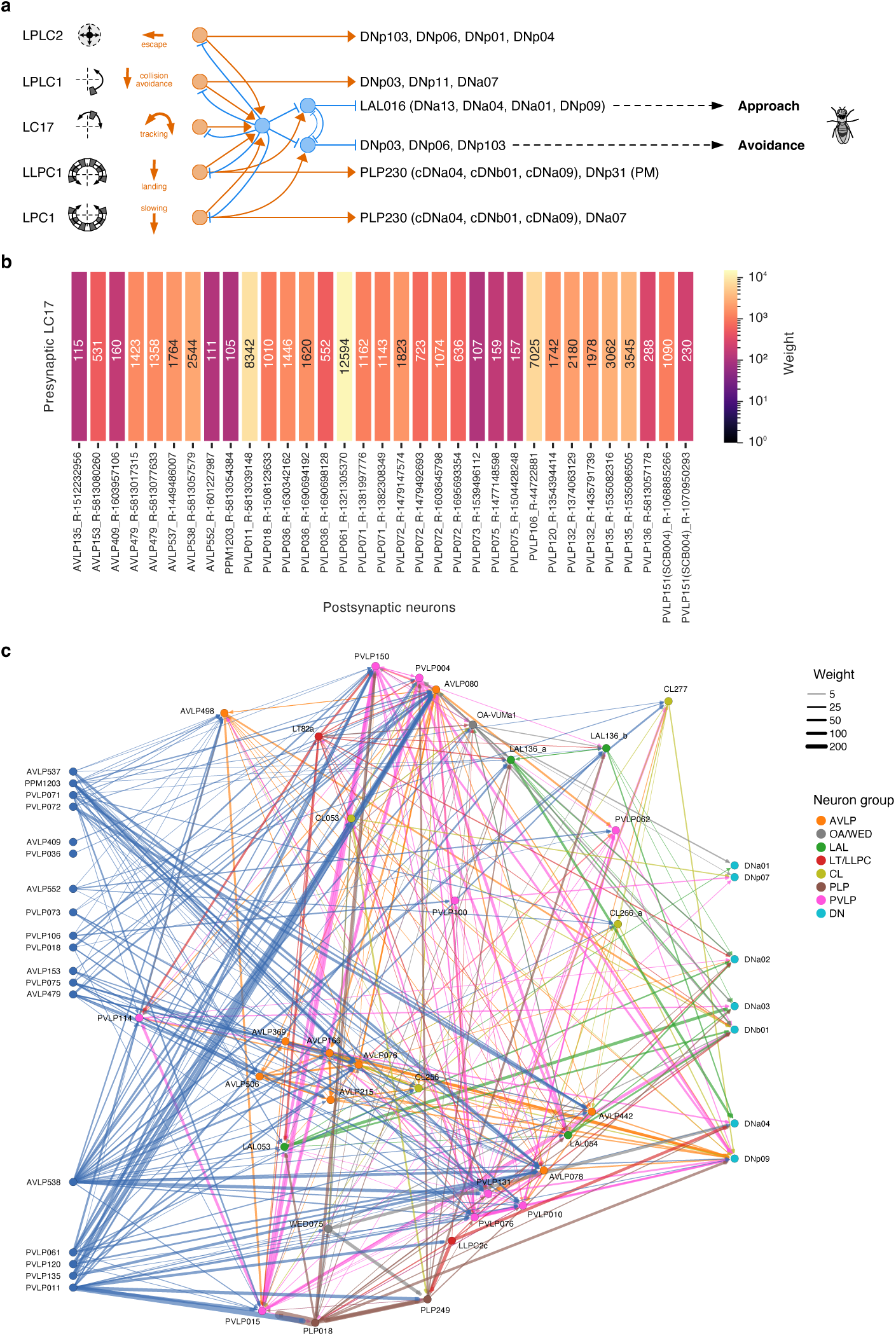
Putative approach circuit downstream LC17 neurons. **a**, Schematic of how different lobula columnar neurons (orange) detecting different visual features connect to a putative GABAergic interneuron (blue on the left, PVLP011) which might be a fundamental hub for approach and avoidance behavior (from FAFB connectome). LC17 is the only type without a mono or disynaptic connection to descending neurons. Two downstream GABAergic interneurons which also receive inputs from translational optic flow detecting neurons (Isaacson, 2019) might release the inhibition for approach (blue on the top-right, PLP249) or avoidance (blue on the bottom-right, PLP018). **b**, LC17 downstream partners from the “hemibrain” connectome. **c**, Pathways connecting the interneurons downstream LC17 with descending neurons (DNs). To note, DNb01 causes ipsilateral decrease and contralateral increase in wing beat amplitude (Ros et al., 2024). PVLP011 is putatively GABAergic and PVLP015 is glutamatergic. The net effect of an LC17 activation would be an ipsilateral increase in DNb01 activity with a consequent ipsilateral turning.

## Methods

### Fly strains

All experiments used 2-5 day old *Drosophila melanogaster* reared on standard cornmeal molasses at 25°C, 30%-50% humidity with a 12 h light – 12 h dark cycle. All behavioral experiments involving manipulations on targeted neurons were performed with flies carrying at least one wild-type *white* allele (aside from the TeTxLC on 1st chromosome). We were not blinded to the genotype tested and mated female flies were randomly chosen from the vials. Fly lines and their origins are listed in Supplementary Table 1.

### Two-photon microscope

Calcium imaging was performed with a VIVO Multiphoton Open (Intelligent Imaging Innovation) system based on a Movable Objective Microscope (MOM, Sutter Instrument) where a mechanical stage (MP-285, Sutter Instrument) allows 3D displacements of the microscope objective. Two-photon excitation was delivered using a Ti:Sapphire laser (Chameleon Vision I, Coherent) tuned to 920 nm with power ranging from 20 to 40 mW (depending on imaging depth). A dual axis galvanometer mirror was used for laser scanning (RGG Scanbox, Sutter Instrument). We imaged with a 20x water-immersion objective (W Plan-Apochromat 20x/1.0 DIC M27, Zeiss) and band-pass filters (Semrock 525/40 and 621/69 nm) were placed in front of the photomultiplier tubes (H11706P-40, Hamamatsu) to reduce background light from the visual display. Microscope and data acquisition were controlled using Slidebook 2024 (Intelligent Imaging Innovation). Single plane images were sampled at ∼9 Hz with a spatial resolution of approximately 180 × 180 pixels (95.7 × 95.7 μm, pixel size ∼0.53 μm, dwell time ∼2 μs). Images and visual stimuli presentation were synchronized a posteriori with frame capture markers (TTL pulses) and stimulus events (analog outputs from the visual display) sampled with a data acquisition device (USB-6229, National Instruments) at 10 kHz.

### 3D-SHOT implementation

Photostimulation via 3D scanless holographic optogenetics with temporal focusing (3D-SHOT^62^) was performed using the Nouveau Phasor (Intelligent Imaging Innovation) module. Two-photon stimulation was enabled by a femtosecond laser (Monaco 1035, Coherent). This technology uses point-cloud computer-generated holography (CGH) combined with temporal focusing (TF) to deliver two-photon spots matching the dimension of a few perikarya (∼10 μm) without limits in their number or z-planes. In brief, CGH uses a spatial light modulator (SLM) to separate the laser beam, after it is expanded, into multiple identical discs of light where the beam intensity determines their dimension, while their different phases determine the 3D holography (through a phase mask displayed on the SLM). This approach alone leads to photostimulation with low spatial resolution along the optical axis (i.e., above and below the focal plane). TF uses a diffraction grating placed before the SLM to decompose the femtosecond pulses into separate wavelengths so that each one propagates with an independent light path. This causes a broadening in time and reduces the density of photons at out-of-focus planes, which in turns reduces the probability of two-photon absorption, leaving a high probability of absorption at the targeted area^104^. The integration of this approach for photostimulation with calcium imaging allows “all-optical” interrogation^63^ (i.e., read and write neural activity) at cellular resolution and millisecond precision across the entire field of view.

### Fly preparation for all-optical interrogation

Flies were head-fixed to a custom fly holder as previously described^35^. Briefly, the fly head was pitched forward pointing down so that the fly eye’s equator held a pitch angle of approximately 60° relative to the imaging plane. The cuticle above the right optic lobe was gently removed and the exposed brain bathed in isotonic saline. Next, the holder was placed under the objective and at the center of a surrounding visual display and calcium responses from targeted neurons recorded with a posterior view.

### Visual stimuli for imaging

A visual display^105^ composed of 48 (8 × 8 dot matrix LEDs) panels arranged in a semi-cylinder was used for visual stimulation as previously described^35^. Four layers of filter (071, LEE Filters) were placed over the display to reduce light intensity (460/243 nm, spectral peak/FWHM). To optimize the extension of the surrounding visual context (±108°, azimuth; ±32°, elevation) and compensate for the fly eye’s angle, the display was tilted forward at 30° from the horizontal plane. The presentation of visual stimuli was controlled using custom-written MATLAB (MathWorks) scripts.

#### Vertical bars experiment

The stimuli were presented to the right half of the display inside a window (72° width × 72° height) ipsilateral to the recording site. Within the window, maximum intensity was used for ON stimuli over a background set to its minimum intensity (i.e., LEDs off), and vice versa for OFF stimuli. Outside the window, the brightness was set to 50% maximum. Stimulus sequence was randomized and repeated 3 times. The duration of each trial was 7.5 s with the first 0.5 s of uniform display illumination (50% maximum intensity), followed by 0.5 s of static window presentation, variable stimulus motion duration (depending on speed, 4.5 s was the longest), and 2 s of static window presentation. Inter-trial time was set to 2 s with the display off to prevent adaptation.

#### Directional sensitivity experiment

A looming stimulus was presented simulating an object approaching the fly at a constant velocity and edges moving in different directions. Specifically, the looming stimulus simulated an object of 0.5 cm radius, starting at a distance of 50 cm and traveling at 62.5 cm/s. The approach caused an expansion from 0.6° to 14° with a loom velocity (l/v) of 8 ms. The edges moved outward from the center of the RF at 20°/s in 24 different directions ranging from 0° to 345° with steps of 15°. Each edge subtended ∼15° at the eye’s equator and swept 15° orthogonal to its orientation, filling a ∼15° square upon completion. The display background was set to 70% maximum intensity while foreground objects (looming or edges) were set to 0% (i.e., LEDs off). The set of visual stimuli was randomly chosen and repeated 2 times. Each trial lasted 4 s and was composed by 0.5 s of uniform background, 0.5 s of visual motion, followed by 3 s of static pattern. Each trial was followed by 3 s of rest in which flies faced the visual background.

### Centering of the receptive field

As previously described^65^, in the directional sensitivity experiment, we identified a responsive neurite from a single neuron to a looming stimulus and then tested its directional sensitivity using moving edges. A custom GUI in MATLAB allowed a real-time scanning for stimulus position. A grid of 48 positions across the right half of the visual display (∼10° per step) was used to identify the center of the RF for a single neuron. Once the RF center was found, the experiment was presented in that specific position on the visual display.

### Data analysis for imaging

Images were exported in .tiff format from Slidebook 2024 and processed following established protocols^35^. We used a MATLAB toolbox developed by Ben J. Hardcastle (available at https://github.com/bjhardcastle/SlidebookObj) to correct for motion artifacts and draw regions of interest (ROI) around individual targeted neurons. The calcium responses were calculated as the average fluorescence intensity of pixels within the ROI (F_t_) at each frame. These mean values were then normalized to a baseline value using the formula: ΔF/F=(F_t_-F_0_)/F_0_ where F_0_ is the mean of F_t_ during the 0.5 s preceding stimulus onset. This approach ensures accurate correction for motion artifacts and reliable quantification of fluorescence intensity changes. Next, the responses for each ROI were then imported in RStudio^106^ using the R package “R.matlab”^107^. Custom R scripts were used for data plotting and statistical analyses.

### Magnetic-tether flight setup

As previously described^66,67^, flies were cold anesthetized and glued to stainless steel pins (0.1 mm diameter, Fine Science Tools). Briefly, pins were trimmed to be 1 cm length and placed on the dorsal thorax in order to get a final fly’s pitch angle of ∼30°. Flies were left to quickly recover upside-down before running the experiment. The visual stimulation was performed using a cylindrical visual display covering 360° in azimuth and 56° in elevation (460/243 nm, spectral peak/FWHM). At the display horizontal midline, each LED subtended an angle of ∼3.75° on the fly’s retina. The fly was suspended between two magnets to allow the animal to freely rotate around its vertical axis (i.e., yaw plane). A circular array of eight infrared diodes (940 nm, emission peak) illuminated the fly from below while an infrared-sensitive camera (Blackfly S USB3, Teledyne FLIR) fitted with a zoom lens (InfiniStix 1.0x/94 mm, Infinity Photo-Optical) and a long pass filter (FGL850M, Thorlabs) recorded the fly’s behavior from the bottom at 200 Hz. To make sure the fly was well tethered, each trial began with 10 s of acclimatation followed by 20 s of wide-field panorama rotating clockwise (CW) and counter-clockwise (CCW) to elicit a strong optomotor response. Flies whose behavior was characterized by excessive wobble were discarded.

### Visual stimuli for behavior

We used a motion-defined bar (30° width × 56° height), a motion-defined small square (15° width × 15° height), and a wide-field panorama (360° width × 56° height) of random bright and dark vertical stripes. These stimuli rotated horizontally around the fly in CW and CCW directions at 112.5°/s. Type of stimulus and speed were chosen based on previous characterizations^31,32,35^. The background pattern was randomly chosen among 96 different options at the beginning of each trial to avoid potential effects of any specific spatial pattern. The stimuli were designed using custom-written MATLAB code. Each trial involved 25 s of rotating stimulus at constant speed and a variable time (5-10 s) of resting period with a static pattern. The object initial position was randomly selected. Each fly was tested twice in six different randomized trials (3 stimuli × 2 directions). The full experiment lasted ∼10 min. Flies that either flew continuously or briefly stopped once throughout the experiment were included in the analysis.

### Optogenetic stimulation for flight behavior

#### Rigid-tether flight setup

Flies were cold anesthetized at ∼4°C and tethered to tungsten pins (0.2 mm diameter) using UV-curable glue (Watch Crystal Glue Clear) as previously described^35^. In brief, the pin was placed on the dorsal thorax orthogonal to the fly’s long axis. The final pitch angle of the fly was between 35° and 45°, which is similar to body angle and wing stroke plane during free hovering^51^. Before running the experiment, flies were left to quickly recover upside-down in a custom designed pin holder at a room temperature of ∼22°C. Next, the fly was positioned above an optical sensor and the wing beat amplitude difference (ΔWBA), which is highly correlated to the fly’s steering effort, was recorded using a DAQ (Digidata 1440A, Molecular Devices) at 1 kHz. Data acquisition was triggered through a voltage step sent by a second DAQ (USB-1208LS, Measurement Computing) interfaced with MATLAB. Randomly, flies expressing CsChrimson tagged with a red fluorescence protein (tdTomato) were dissected after the behavioral experiment and genetic expression confirmed with a fluorescence stereomicroscope (SteREO Discovery.V12, Zeiss).

#### Unilateral stimulation

Optogenetic activations in behavioral experiments were performed using a fiber-coupled red LED (M660FP1, Thorlabs; 660/20 nm, spectral peak/FWHM) fitted with a zoom lens (MAP1075150, Thorlabs) positioned posteriorly to the tethered fly. The narrow beam was focused over the right optic lobe (∼300 μm in diameter). All-trans-retinal (ATR) is required to get a proper CsChrimson protein conformation. In order to boost flies’ performance, although flies endogenously produce retinal, we added ATR to the food. The progeny from crosses between Gal4 driver lines and UAS-CsChrimson were raised in the darkness to avoid channels’ stimulation. After eclosion, newborn flies were transferred in 0.5 mM ATR food and kept there for 3-5 days until the experiment.

### Data analysis for flight behavior

Data collected in both rigid-tether and magnetic-tether setups were analyzed in RStudio^106^ using custom R scripts. Video recordings from magnetic-tether experiments were imported in MATLAB and the flies’ heading extracted using a custom software. Body saccades were detected as previously described^31^. Data plotting was performed using R package “ggplot2”^108^.

### Optogenetic stimulation for walking behavior

#### Rearing

Experimental flies were collected over three consecutive days following the first eclosion from a cross, and then sorted (selection against balancers) on a cold plate. Flies were subsequently housed in low-density vials (6-8 flies per vial) on regular food for 2-4 days. The day prior to experiments, flies were moved to starvation vials, which consisted of a Kimwipe (Kimberly-Clark) saturated with 2 mL of a 0.5 mM ATR dilution solution. Before commencing experiments, flies were tethered with a pin and allowed to acclimate for 5-10 min.

#### Walking setup

The experimental setup utilized a custom-designed fly-on-a-ball arena, enclosed by black acrylic panels and covered with black cloth to ensure complete darkness. A custom-designed holder air-suspended the ball (8 mm in diameter), maintaining a constant flow of humidified air at 300 mL/min. Ball movement was recorded and tracked using fictrac^109^ via a Raspberry Pi camera (ArduCam) positioned behind the fly. The fly was tethered vertically onto the ball at an angle of approximately 15° to encourage walking. Two fiber optic light sources (Adafruit; 625 nm, spectral peak) were diagonally positioned and focused on the fly’s head using thin, rigid fiber optic cables. The ball was tracked at 80 Hz, and experimental timekeeping was performed at 60 Hz. All experimental parameters were controlled and recorded using python scripts on Robot Operating System (ROS). Data analysis was conducted using Python.

### Generation of smGdP-10XV5-tagged allele of *shakB*

Constitutive and conditional alleles of *shakB* were generated as previously described^84^. A Spaghetti Monster V5 tag (smGdP-10xV5) was inserted at the C-terminus immediately upstream of the stop codon, thereby labeling all annotated isoforms in FlyBase (FB2025_03). Primer sequences and plasmid maps are available upon request.

### Immunohistochemistry, tissue clearing and DPX mounting

All protocols in immunohistochemistry and DPX (dibutyl phthalate in xylene) mounting were performed as described in a previous study^8^.

### Antibody information

#### Primary antibodies and dilutions used in this study

Chicken anti-GFP (1:1000, Abcam #ab13970, RRID: AB_300798), rabbit anti-dsRed (1:200, Clontech #632496, RRID: AB_10013483), mouse anti-Bruchpilot (1:20, DSHB Nc82, RRID: AB_2314866), chicken anti-V5 (1:200, Fortis Life Sciences #A190-118A, RRID: AB_66741), mouse anti-V5 (1:500, Abcam #ab27671, RRID: AB_471093), rabbit anti-HA (1:200, Cell Signaling Technology #3724, RRID: AB_1549585), rat anti-N-Cadherin (1:40, DSHB MNCD2, RRID: AB_528119), anti-GFP nanobody (1:200 for expansion microscopy, NanoTag Biotechnologies #N0304-At488-L, RRID: AB_2744629).

#### Secondary antibodies and dilutions used in this study

Goat anti-chicken AF488 (1:500, Invitrogen #A11039, RRID: AB_2534096), goat anti-mouse IgG2A (1:500, Invitrogen #A21131, RRID: AB_2535771), goat anti-rabbit AF568 (1:500, Invitrogen #A11011, RRID: AB_143157), goat anti-mouse AF647 (1:500, Jackson ImmunoResearch #115-607-003, RRID: AB_2338931), goat anti-rat AF647 (1:500, Jackson ImmunoResearch #112-605-167, RRID: AB_2338404).

### Confocal image acquisition and processing

Immunofluorescence images were acquired using a confocal microscope (LSM 880, Zeiss) equipped with an oil-immersion objective (Plan-Apochromat 63x/1.4 Oil DIC M27, Zeiss) and the digital imaging software ZEN (Zeiss) which controlled the 488 nm, 561 nm, and 633 nm lasers. Serial optical sections were obtained from whole-mount brains with a typical resolution of 1024 μm × 1024 μm, and 0.5 μm thick optical sections. Image stacks were then imported to either Fiji 2.0.0-rc-69/1.52k^110^ or Imaris 10.1 (Oxford Instruments) for level adjustment, cropping, removal of off-target brain regions and background noise, and 3D volume reconstructions.

### Expansion Light Sheet Microscopy (ExLSM) and HCR-FISH

Tissue staining, gelation and expansion for ExLSM protocols were adapted from a previous work^65^. After dissection, fixation and permeabilization, brains were stored in RNAse-free 0.5% PBST containing anti-GFP nanobody (NanoTag Biotechnologies #N0304-At488-L) overnight at 4°C. All samples were subsequently processed using a protein and RNA retention ExM protocol with minor modifications^111^ and adjustments for the fly brain^65^.

#### HCR-FISH

Following digestion with proteinase K, gels with embedded brains were washed 3 times with PBS, transferred into 24-well plates (Laguna Scientific #4624-24), and digested with DNAse diluted in RDD buffer (RNase-Free DNase Set, Qiagen #79254) to limit DAPI signal to RNA only and facilitate subsequent analysis for 2 h at 37°C. After three washes in PBS, gels were equilibrated in Probe Hybridization Buffer (Molecular Instruments) for 30 min at 37°C, and then transferred to new 24-well plates containing custom-designed probes (Molecular Instruments) diluted in pre-warmed Probe Hybridization Buffer (1 μL of 1 μM stock probe solution per 200 μL of buffer) and left shaking overnight at 37°C. The following day, gels were washed 4 times with pre-warmed Probe Wash Buffer (Molecular Instruments) for 20 min at 37°C, then washed twice for 5 min with SSCT buffer (SSC, Thermo Fisher #AM9763 with 0.05% Triton X-100) at room temperature and transferred to new 24-well plates with HCR Amplification buffer (Molecular Instruments) for equilibration. Hairpins (HCR Amplifiers, Molecular Instruments) conjugated with AF546 dye were diluted in Amplification Buffer (2 μL of each hairpin per 100 μL of buffer), heat-activated in a thermal cycler (90 s at 95°C), removed, and kept for 30 min at room temperature in the dark. After 30 min, the hairpins were added to the 24-well plates with gels (300 μL per well) and incubated with shaking at room temperature in the dark for 3 h. The hairpin solution was then removed, and the gels were washed 4 times with SSCT and 2 times with SSC for 10 min at RT in the dark. Gels were subsequently stored at 4°C in SSC until final expansion.

#### Sample mounting

Samples were expanded to approximately 3x in 0.5x PBS containing 1:1,000 SYTO-DAPI (Thermo Fisher S11352) at room temperature for 2 h before mounting onto poly-l-lysine (PLL)-coated coverslips. The coverslips were then bonded with Bondic UV-curing adhesive (Bondic starter kit, Bondic) onto a custom-fabricated sample holder (Janelia Tech ID 2021-021) to be suspended vertically in the imaging chamber. Mounted samples were imaged in 0.5x PBS with 1:10,000 SYTO-DAPI after a minimum of 1 h of equilibration in the imaging chamber. Unexpanded gels were stored at 4°C in 1x PBS + 0.02% sodium azide (Sigma-Aldrich, Cat# S8032) for up to 14 days before final expansion and imaging.

#### Light sheet microscopy

Images were acquired on a microscope (LS7, Zeiss) equipped with 405 nm, 488 nm, 561 nm, and 638 nm lasers. Illumination optics with a 10x/0.2 NA were used for excitation (Zeiss, Cat# 400900-9020-000). Detection was performed using a water-immersion objective (W Plan-Apochromat 20x/1.0 DIC M27, Zeiss Cat# 421452-9700-000). The microscope optical zoom was set to 2.5x, resulting in a total magnification of 50x. DAPI and AF546 dyes were simultaneously excited by the 405 nm and 561 nm laser lines, and emission light was separated by a dichroic mirror (SBS LP 510, Zeiss Cat# 404900-9312-000) with a band-pass filter 445/50 nm (spectral peak/FWHM) and a modified filter 527/23 nm (Chroma, Cat# ET672/23m). Similarly, AF488 and SeTau647 dyes were simultaneously excited via 488 nm and 638 nm, and the emission was split through a dichroic mirror (SBS LP 560, Zeiss) with a band-pass filter 525/40 nm and long-pass 660 nm (Zeiss Cat# 404900-9318-000). To eliminate laser transmission, a 405/488/561/640 nm laser blocking notch filter (Zeiss, Cat# 404900-9101-000) was added to the emission path. Images were captured using dual PCO.edge 4.2 detection modules (Zeiss, Cat# 400100-9060-000) with a 50 ms exposure time. Filter and camera alignment were manually calibrated prior to each imaging session. Image volumes were acquired at optimal z-step and light-sheet thickness, and the Pivot Scan feature was used to reduce illumination artifacts by sweeping the light-sheet in the x-y plane. The microscope was operated using ZEN Black 3.1 (v9.3.6.393, Zeiss).

### Analysis of HCR-FISH data from ExLSM Image Stacks

The full details of our analysis are available in our previous publications^65,112^. The acquired light sheet z-stacks, stored in CZI format, were imported and pre-processed to remove noise and artifacts generated by the imaging modality. These artifacts included limited channel contrast, variations in contrast across images within a dataset, background noise fluctuations due to both intra-channel variations and inter-channel crosstalk, and localized brightness changes caused by varying fluorophore concentrations within and among stained nuclei. Pre-processing involved the following steps: full-scale contrast stretching (FSCS) to normalize luminosity across different channels, local background removal using a 3D Gaussian filter, a second FSCS to compensate for any contrast loss due to background removal, and a final median filter to eliminate any remaining localized noise. These pre-processed stacks served as the starting point for instance segmentation of the nuclei. First, nuclear centers were identified using a Laplacian-of-Gaussian (LoG) filter. Then, the imaged volume was subdivided into 3D Voronoi cells, using the detected centers as seeds and Euclidean distance. Each cell contained one nucleus, which was segmented using a threshold obtained by minimizing an energy functional designed to find the optimal surface separating the nucleus from the surrounding cytoplasm. Once nuclei were segmented, the FISH puncta were identified within the associated 3D volume using a LoG filter, and only the puncta within the nucleus region and its immediate surrounding volume were counted. Pre-processed products, segmented nuclear features, associated FISH puncta, and their features were stored for further analysis. The package was written in Python, is open-source, and is available for download at https://github.com/avaccari/DrosophilaFISH.

### Intracellular neurobiotin filling

#### Whole-cell patch-clamp

Electrophysiological recordings with neurobiotin filling were performed in flies expressing mCD8-GFP and RNAi for *Fmr1* in LC17 neurons. The GFP was used to visualize the neurons of interest, while the knockdown of *Fmr1* was done to increase neurobiotin uptake into the neurons and improve the visualization of electrically coupled circuits^100^. Adult male or female flies (2-3 days old) were anesthetized on ice and their brain dissected on a Sylgard-coated lid of a Terasaki plate in saline containing 103 mM NaCl, 3 mM KCl, 1.5 mM CaCl_2_, 4 mM MgCl_2_, 26 mM NaHCO_3_, 1 mM NaH_2_PO_4_, 5 mM N-Tris (hydroxymethyl) methyl-2-aminoethane-sulfonic acid (TES), 10 mM trehalose, 10 mM glucose, 2 mM sucrose (adjusted to 273-275 mOsm, pH equilibrated to 7.3-7.4)^113^. Each brain was then transferred onto a PLL-coated custom-cut coverslip (3 mm × 3 mm) to keep it in place during the recording. The brains were positioned with a frontal view (i.e., facing up), the glial sheath removed around LC17 somata using fine forceps (Dumont #5SF, Fine Science Tools), and the optic lobes gently stretched to better expose the cell bodies in the ventral region. Next, the coverslip with the prepared brain was placed under a direct microscope (Axioplan 2, Zeiss), equipped with a water-immersion objective (LUMPlanFL/IR 60x/0.9 W, Olympus), for patching. Whole-cell patch-clamp recordings from LC17 were obtained with saline continuously perfused (2 mL/min) and bubbled (95% O_2_/5% CO_2_) over the preparation. A borosilicate glass pipette with tip-resistance between 10 and 15 MΩ was filled with internal solution containing 140 mM potassium aspartate, 10 mM HEPES, 1 mM KCl, 4 mM MgATP, 0.5 mM Na_3_GTP, 1 mM EGTA, 10 mM neurobiotin (SP-1120, Vector Laboratories), 20 μM Alexa Fluor 594-hydrazide-Na (260-275 mOsm, pH 7.3). Only if a seal resistance >5 GΩ before breaking in, and a stable recording >15 min were achieved, was the brain processed for subsequent staining. Recordings were acquired in current clamp mode with a MultiClamp 700B (Molecular Devices) amplifier, low-pass filtered at 4 kHz, and digitalized at 10 kHz with Digidata 1300b (Molecular Devices). A series of step current injections in bridge mode were applied to the cell to elicit action potentials and iontophoretically injected neurobiotin using the software pCLAMP (Molecular Devices). Only one cell per optic lobe was patched and filled. After the injection, neurobiotin was allowed to diffuse via gap junctions for ∼30 min.

#### Immunohistochemistry

Following the fillings, the brains were fixed with 4% v/v PFA (158127, Sigma-Aldrich) in PBS for 30 min at room temperature. Next, the fixative was washed out with PBS, the brains solubilized in PBST (0.5% Triton X100, T9284, Sigma-Aldrich) for 1 h, and blocked in 7.5% v/v NGS (Normal Goat Serum, 005-000-121, Jackson ImmunoResearch) in PBST for 1-2 h at room temperature. Brains were then incubated in primary antibodies: chicken anti-GFP (1:1000, Abcam #ab13970), mouse anti-Bruchpilot (1:10, DSHB nc82, RRID: AB_2314866), and streptavidin-AF647 (1:500, Invitrogen S21374) for 4 h at room temperature and 24 h at 4°C. Samples were subsequently washed out of primary antibodies at least 3 times with PBST over 2 h and incubated in secondary antibodies: goat anti-chicken AF488 (1:100, Invitrogen #A11039, RRID: AB_2534096) and goat anti-mouse AF568 (1:200, Invitrogen #A11004, RRID: AB_2534072) for 4 h at room temperature and for 48 h at 4°C. After washing the samples at least 3 times with PBST over 2 h, they were finally incubated in VectaShield (H-1000, Vector Laboratories) for 10 min at room temperature, and mounted onto slides for imaging. Brains were imaged with a confocal microscope (see description for confocal image acquisition above). Complete series of optical sections were taken at 1 μm intervals with an oil-immersion objective (Plan-Apochromat 40x/1.3 Oil DIC M27, Zeiss).

### scRNA-seq analysis

The expression levels of *shakB* were obtained from the scRNA-seq atlas of the developing *Drosophila* visual system^25,81^. The data were analyzed and plotted using RStudio^106^.

### Connectomics analysis of the optic lobe

Two electron microscopy reconstructed volumes (FAFB^21^ and MAOL^22^) were explored using the R package “coconatfly”^59^ which builds upon the R package “natverse”^114^. The neurons of interest were loaded and their pre- or post-synaptic partners retrieved without setting any threshold (i.e., every synapse). Only the right optic lobe was considered. For clarity, the pie charts grouped together partners with less than 1000 or 3000 synapses (depending on the dataset). This allowed us to highlight and visualize only the most influential partners. We noticed that the number of synapses were in general much higher for MAOL than FAFB but the relative connection strengths to partners maintain a sufficient consistency between the two datasets. This permitted us to confidently ascribe the difference to variability in synapse detection algorithms or proofreading. We also matched cell types across the two datasets finding strong overlap with a previous comprehensive and systematic matching of all the optic lobe neurons^22^.

### Connectomics analysis of pathways to descending neurons

We explored the “hemibrain” connectome^49^ to find pathways from LC17 to descending neurons using the python API “neuPrint”^115^ and custom code. No direct connections between LC17 neurons to descending neurons were identified in the hemibrain connectome. To systematically identify pathways linking the LC17 pathway to descending neurons, we first identified the set of neurons that were common downstream partners of LC17 neurons and that received at least 100 synapses from the LC17 population. This yielded a set of 33 interneurons, which were used as the source population for subsequent analysis. We then defined the complete set of descending neurons as the target population and enumerated all directed pathways of three hops or fewer between each source neuron–descending neuron pair where each edge has at least 10 synapses. All pathways that route back through LC17 were discarded. For visualization, we applied an additional filtering step to simplify the network. First, we only display selected descending neuron populations (DNa01, DNa02, DNa03, DNa04, DNa10, DNb01, DNp07, and DNp09). Next, we retained only two-hop (or fewer) pathways containing a single intermediate neuron (source neuron → intermediate neuron → descending neuron), in which both constituent edges had synaptic weights greater than 10. We then collected all neurons participating in these filtered two-hop pathways and extracted the complete subnetwork among them. Consequently, the displayed network contains not only the filtered two-hop source-to-descending motifs, but all synaptic connections among the selected neurons, including edges that participate in shorter or longer pathways within the subnetwork. For graph layout, the LC17 common downstream partner neurons were positioned along one side of the network and descending neurons along the opposite side. All nodes represent neuron populations (grouping individual neurons of the same type into one node). A force-directed spring layout was then applied to the full graph using synaptic weight as the edge-weight parameter, allowing intermediate nodes to settle according to their weighted connectivity while preserving the fixed anchor positions of the source and target populations. After layout optimization, the x-coordinates of the intermediate nodes were rescaled to a predefined horizontal range spanning the space between the anchored node classes. The vertical ordering of nodes (across all layers) was determined by hierarchical clustering based on connectivity similarity (method = “ward”, metric = “euclidean”) within the subnetwork.

### Statistics

Generalized linear models (GLM) were fitted to the data using built-in R functions. Linear models with Gaussian errors fitted normally distributed data, whereas GLM with Gamma errors and a log link function fitted strongly skewed positive data. Mixed effects models were also used when the data were not averaged per fly, preserving statistical power and allowing to adjust estimates for repeated sampling or sample unbalancing. In this later case, the R package “lme4”^116^ was used and the individual flies were chosen as random effects. Pairwise post-hoc comparisons (*t*-test) on the estimates of the model were performed using the R package “emmeans”^117^. Using the same package, we calculated the Cohen’s d effect sizes as the pairwise difference between model estimates divided by the standard deviation of the data. Dara plotting was performed with the R package “ggplot2”^108^. In violin and box plots, both mean and median were reported as central tendency measures, bottom and top edges of the box represent 25th (Q1) and 75th (Q3) percentiles and whiskers represent the lowest and highest datum within 1.5 interquartile range (Q3 - Q1). No power analysis was performed to determine sample size. Based on the literature in the field, we determined the sample size for each specific experiment.

## Acknowledgements

We thank Georg Ammer for sharing the *shakB*-Trojan-Gal4 gene trap line; Mert Erginkaya and Jan M. Ache for preliminary data; Orkun Akin and his lab members for advice on fly preparation for electrophysiology, and for sharing lab space and equipment; Margot Quinlan for sharing reagents; Jeffrey M. Donlea and his lab members for sharing lab space and equipment; Bloomington *Drosophila* Stock Center (NIH P40OD018537) and Janelia Fly Facility for providing fly strains; members of the FlyWire Consortium for connectome proofreading and annotation to the FAFB dataset (supported by NIH BRAIN Initiative RF1MH117815, RF1MH129268 and U24NS126935); developers of the natverse including the coconatfly and fafbseg packages (supported by NIH BRAIN Initiative 1RF1MH120679, NSF/MRC Neuronex2 NSF 2014862/MC_EX_MR/T046279/1, and core funding from the Medical Research Council MC-U105188491). This work was supported by NEI K99EY036889 to G.F.; K99EY036123 to M.D; NINDS R01NS054814 to V.H.; and NEI R01EY026031 to M.A.F.

## Authors contributions

G.F. and M.A.F. conceived and designed the study. G.F. performed and analyzed the data of behavioral and calcium imaging experiments. G.F. performed connectome analysis. M.D., Parmis S.M., Pegah S.M., and P.S. conducted molecular genetics and immunohistochemical experiments, with S.L.Z assisting with data interpretation. M.D. and Parmis S.M., designed and performed HCR-FISH experiments, with A.V. analyzing the data. Y.Z.K. analyzed scRNA-seq datasets. P.K. performed behavioral experiments in walking flies, with V.H. assisting with data interpretation. P.M. conducted electrophysiological experiments, with G.F. and L.M.P.C. assisting with brain preparation. G.F., M.D. and M.A.F. wrote the manuscript, with substantial input and feedback from S.L.Z. All authors reviewed and edited the manuscript.

