## Supplemental Table 1 for "Electrical synapses mediate visual approach behavior"

| Figure | Cells targeted | Description in Figure | Chr | Fly Genotype | Source |
|---|---|---|---|---|---|
| 1f-g | Control | All-optical interrogations | 1st | 13xLexAop-IVS-jGCaMP7f / w[1118] | BDSC #80910 |
| Ext. Fig. 1c | LC17 | | 2nd | R21D03-LexA / empty-p65.AD | BDSC #52590 |
| | | | 3rd | empty-GAL4.DBD / 20xUAS-IVS-Syn21-Chrimson::tdTomato-3.1 (VK00005) | BDSC #79603 (empty split-GAL4) / D. Anderson Lab |
| 1f-i | T3 | All-optical interrogations | 1st | 13xLexAop-IVS-jGCaMP7f / w[1118] | BDSC #80910 |
| Ext. Fig. 1c | LC17 | | 2nd | R21D03-LexA / VT002055-p65.AD | BDSC #52590 / BDSC #73391 |
| | | | 3rd | R65B04-GAL4.DBD / 20xUAS-IVS-Syn21-Chrimson::tdTomato-3.1 (VK00005) | BDSC #69453 (Keleş et al., 2020) / D. Anderson Lab |
| Ext. Fig. 1d,e | T3 | All-optical interrogations | 1st | 13xLexAop2-Syn21-opGCaMP6s (su(Hw)attP8), 10xUAS-Syn21-Chrimson88::tdTomato-3.1 (attP18) / w[1118] | M. Reiser Lab (Strother et al., 2017) |
| | LC17 | | 2nd | R21D03-LexA / VT002055-p65.AD | BDSC #52590 / BDSC #73391 |
| | | | 3rd | R65B04-GAL4.DBD / + | BDSC #69453 (Keleş et al., 2020) |
| Ext. Fig. 1f-h | T3 | All-optical interrogations | 1st | 13xLexAop-IVS-jGCaMP7f / w[1118] | BDSC #80910 |
| | LC17 | | 2nd | R21D03-LexA / VT002055-p65.AD | BDSC #52590 / BDSC #73391 |
| | | | 3rd | R65B04-GAL4.DBD / 20xUAS-Chronos::mVenus | BDSC #69453 (Keleş et al., 2020) / BDSC #77117 |
| Ext. Fig. 2a | Control | Rigid tether flight behavior | 1st | w[1118] / w[1118] | |
| | | Chrimson - Optogenetic | 2nd | empty-p65.AD / + | |
| | | | 3rd | empty-GAL4.DBD / 20xUAS-IVS-Syn21-Chrimson::tdTomato-3.1 (VK00005) | BDSC #79603 (empty split-GAL4) / D. Anderson Lab |
| Ext. Fig. 2a | LC17 | Rigid tether flight behavior | 1st | w[1118] / w[1118] | |
| | | Chrimson - Optogenetic | 2nd | R21D03-p65.AD / + | |
| | | | 3rd | R65C12-GAL4.DBD / 20xUAS-IVS-Syn21-Chrimson::tdTomato-3.1 (VK00005) | BDSC #68356 (OL0005B split-GAL4) / D. Anderson Lab |
| Ext. Fig. 2c | Control | Rigid tether walking behavior | 1st | w[1118] / w[1118](31 + CS)6x | |
| | | CsChrimson - Optogenetic | 2nd | empty-p65.AD / 20xUAS-IVS-CsChrimson::mVenus | BDSC #55135 |
| | | | 3rd | empty-GAL4.DBD / + | M. Dickinson Lab (empty split-GAL4) |
| Ext. Fig. 2c | LC17 | Rigid tether walking behavior | 1st | w[1118] / w[1118](31 + CS)6x | |
| | | CsChrimson - Optogenetic | 2nd | R21D03-p65.AD / 20xUAS-IVS-CsChrimson::mVenus | BDSC #55135 |
| | | | 3rd | R65C12-GAL4.DBD / + | BDSC #68356 (OL0005B split-GAL4) |
| 2b-g | LC11 | Calcium imaging | 1st | w[1118] / w[1118] | |
| Ext. Fig. 3b,c | | | 2nd | R22H02-p65.AD / + | |
| | | | 3rd | R20G06-GAL4.DBD / 20xUAS-IVS-jGaCMP7f | BDSC #68362 (OL0015B split-GAL4) / BDSC #79031 |
| 2b-i | LC17 | Calcium imaging | 1st | w[1118] / w[1118] | |
| Ext. Fig. 3a-c | | | 2nd | R21D03-p65.AD / + | |
| | | | 3rd | R65C12-GAL4.DBD / 20xUAS-IVS-jGaCMP7f | BDSC #68356 (OL0005B split-GAL4) / BDSC #79031 |
| 2h-i | LPLC2 | Calcium imaging | 1st | UAS-Dcr2 / w[1118] | BDSC #24646 |
| | | | 2nd | UAS-Control RNAi / + | VDRC #60100 |
| | | | 3rd | VT049479-GAL4 / 20xUAS-IVS-jGCaMP7f | Janelia Fly Facility, BDSC #79031 |
| 3c-e | Control | Magnetic tether flight behavior | 1st | w[+] / w[1118] | |
| Ext. Fig. 4b | | Kir2.1 - Silcencing | 2nd | empty-p65.AD / + | |
| 5d,e - Ext. Fig. 8 | | | 3rd | empty-GAL4.DBD / 10xUAS-IVS-eGFP::Kir2.1 | BDSC #79603 (empty split-GAL4) / C. von Reyn Lab |
| 3c-e | LC17 | Magnetic tether flight behavior | 1st | w[+] / w[1118] | |
| Ext. Fig. 4a,b | | Kir2.1 - Silcencing | 2nd | R21D03-p65.AD / + | |
| 5d,e - Ext. Fig. 8 | | | 3rd | R65C12-GAL4.DBD / 10xUAS-IVS-eGFP::Kir2.1 | BDSC #68356 (OL0005B split-GAL4) / C. von Reyn Lab |
| 3c-e | T3 | Magnetic tether flight behavior | 1st | w[+] / w[1118] | |
| Ext. Fig. 4a,b | | Kir2.1 - Silcencing | 2nd | VT002055-p65.AD / + | BDSC #73391 |
| | | | 3rd | R65B04-GAL4.DBD / 10xUAS-IVS-eGFP::Kir2.1 | BDSC #69453 (Keleş et al., 2020) / C. von Reyn Lab |
| 3c-e | T4/T5 | Magnetic tether flight behavior | 1st | w[+] / w[1118] | |
| Ext. Fig. 4b | | Kir2.1 - Silcencing | 2nd | R59E08-p65.AD / + | |
| | | | 3rd | R42F06-GAL4.DBD / 10xUAS-IVS-eGFP::Kir2.1 | BDSC #86858 (SS00324 split-GAL4) / C. von Reyn Lab |
| 3c-e | T2a | Magnetic tether flight behavior | 1st | w[+] / w[1118] | |
| Ext. Fig. 4b | | Kir2.1 - Silcencing | 2nd | VT013616-p65.AD / + | |
| | | | 3rd | VT025776-GAL4.DBD / 10xUAS-IVS-eGFP::Kir2.1 | BDSC #88945 (SS02353 split-GAL4) / C. von Reyn Lab |
| 3f-h | Control | Magnetic tether flight behavior | 1st | UAS-TeTxLC.tntC1 / w[1118] | BDSC #28996 |
| Ext. Fig. 4c,f | | TeTxLC (X) - Synaptic blockade | 2nd | empty-p65.AD / + | |
| 5d,e - Ext. Fig. 8 | | | 3rd | empty-GAL4.DBD / + | BDSC #79603 (empty split-GAL4) |
| 3f-h | LC17 | Magnetic tether flight behavior | 1st | UAS-TeTxLC.tntC1 / w[1118] | BDSC #28996 |
| Ext. Fig. 4c,f | | TeTxLC (X) - Synaptic blockade | 2nd | R21D03-p65.AD / + | |
| 5d,e - Ext. Fig. 8 | | | 3rd | R65C12-GAL4.DBD / + | BDSC #68356 (OL0005B split-GAL4) |
| 3f-h | T3 | Magnetic tether flight behavior | 1st | UAS-TeTxLC.tntC1 / w[1118] | BDSC #28996 |
| Ext. Fig. 4c,f | | TeTxLC (X) - Synaptic blockade | 2nd | VT002055-p65.AD / + | BDSC #73391 |
| | | | 3rd | R65B04-GAL4.DBD / + | BDSC #69453 (Keleş et al., 2020) |
| 3f-h | T4/T5 | Magnetic tether flight behavior | 1st | UAS-TeTxLC.tntC1 / w[1118] | BDSC #28996 |
| Ext. Fig. 4c | | TeTxLC (X) - Synaptic blockade | 2nd | R59E08-p65.AD / + | |
| | | | 3rd | R42F06-GAL4.DBD / + | BDSC #86858 (SS00324 split-GAL4) |
| 3f-h | T2a | Magnetic tether flight behavior | 1st | UAS-TeTxLC.tntC1 / w[1118] | BDSC #28996 |
| Ext. Fig. 4c,f | | TeTxLC (X) - Synaptic blockade | 2nd | VT013616-p65.AD / + | |
| | | | 3rd | VT025776-GAL4.DBD / + | BDSC #88945 (SS02353 split-GAL4) |
| 3i-k | Control | Magnetic tether flight behavior | 1st | w[+] / w[1118] | |
| Ext. Fig. 4d | | shakB[RNAi] - Gap junction knockdown | 2nd | empty-p65.AD / + | |
| 5d,e - Ext. Fig. 8 | | | 3rd | empty-GAL4.DBD / UAS-shakB[RNAi] (VALIUM20) | BDSC #79603 (empty split-GAL4) / BDSC #57706 |
| 3i-k | LC17 | Magnetic tether flight behavior | 1st | w[+] / w[1118] | |
| Ext. Fig. 4d | | shakB[RNAi] - Gap junction knockdown | 2nd | R21D03-p65.AD / + | |

| | | | | | |
|---|---|---|---|---|---|
| 5d,e - Ext. Fig. 8 | | | 3rd | R65C12-GAL4.DBD / UAS-shakB[RNAi] (VALIUM20) | BDSC #68356 (OL0005B split-GAL4) / BDSC #57706 |
| 3i-k | T3 | Magnetic tether flight behavior | 1st | w[+] / w[1118] | |
| Ext. Fig. 4d | | shakB[RNAi] - Gap junction knockdown | 2nd | VT002055-p65.AD / + | BDSC #73391 |
| | | | 3rd | R65B04-GAL4.DBD / UAS-shakB[RNAi] (VALIUM20) | BDSC #69453 (Keleş et al., 2020) / BDSC #57706 |
| 3i-k | T4/T5 | Magnetic tether flight behavior | 1st | w[+] / w[1118] | |
| Ext. Fig. 4d | | shakB[RNAi] - Gap junction knockdown | 2nd | R59E08-p65.AD / + | |
| | | | 3rd | R42F06-GAL4.DBD / UAS-shakB[RNAi] (VALIUM20) | BDSC #86858 (SS00324 split-GAL4) / BDSC #57706 |
| 3i-k | T2a | Magnetic tether flight behavior | 1st | w[+] / w[1118] | |
| Ext. Fig. 4d | | shakB[RNAi] - Gap junction knockdown | 2nd | VT013616-p65.AD / + | |
| | | | 3rd | VT025776-GAL4.DBD / UAS-shakB[RNAi] (VALIUM20) | BDSC #88945 (SS02353 split-GAL4) / BDSC #57706 |
| Ext. Fig. 4e | Pan-neuronal | Magnetic tether flight behavior | 1st | w[+] / w[1118] | |
| | | shakB[RNAi] - Gap junction knockdown | 2nd | + / + | |
| | | | 3rd | R57C10-GAL4 / UAS-shakB[RNAi] (VALIUM20) | BDSC #39171 / BDSC #57706 |
| Ext. Fig. 4g | Control | Magnetic tether flight behavior | 1st | w[+] / w[1118] | |
| | | TeTxLC (II) - Synaptic blockade | 2nd | empty-p65.AD / UAS-TeTxLC.tntG2 | BDSC #28838 |
| | | | 3rd | empty-GAL4.DBD / + | BDSC #79603 (empty split-GAL4) |
| Ext. Fig. 4g | LC17 | Magnetic tether flight behavior | 1st | w[+] / w[1118] | |
| | | TeTxLC (II) - Synaptic blockade | 2nd | R21D03-p65.AD / UAS-TeTxLC.tntG2 | BDSC #28838 |
| | | | 3rd | R65C12-GAL4.DBD / + | BDSC #68356 (OL0005B split-GAL4) |
| Ext. Fig. 4g | T3 | Magnetic tether flight behavior | 1st | w[+] / w[1118] | |
| | | TeTxLC (II) - Synaptic blockade | 2nd | VT002055-p65.AD / UAS-TeTxLC.tntG2 | BDSC #73391 / BDSC #28838 |
| | | | 3rd | R65B04-GAL4.DBD / + | BDSC #69453 (Keleş et al., 2020) |
| Ext. Fig. 4g | T4/T5 | Magnetic tether flight behavior | 1st | w[+] / w[1118] | |
| | | TeTxLC (II) - Synaptic blockade | 2nd | R59E08-p65.AD / UAS-TeTxLC.tntG2 | BDSC #28838 |
| | | | 3rd | R42F06-GAL4.DBD / + | BDSC #86858 (SS00324 split-GAL4) |
| Ext. Fig. 4g | T2a | Magnetic tether flight behavior | 1st | w[+] / w[1118] | |
| | | TeTxLC (II) - Synaptic blockade | 2nd | VT013616-p65.AD / UAS-TeTxLC.tntG2 | BDSC #28838 |
| | | | 3rd | VT025776-GAL4.DBD / + | BDSC #88945 (SS02353 split-GAL4) |
| Ext. Fig. 5a,b | Control | Expressing nuclear GFP | 1st | w[1118] / w[1118] | |
| | | HCR-FISH | 2nd | R21D03-p65.AD / UAS-His2A-GFP | L. Zipursky Lab |
| | | mCherry[RNAi] - Control knockdown | 3rd | R65C12-GAL4.DBD / UAS-mCherry[RNAi] (VALIUM20) | BDSC #68356 (OL0005B split-GAL4) / BDSC #35785 |
| Ext. Fig. 5a,b | LC17 | Expressing nuclear GFP | 1st | w[1118] / w[1118] | |
| | | HCR-FISH | 2nd | R21D03-p65.AD / UAS-His2A-GFP | L. Zipursky Lab |
| | | shakB[RNAi] - Gap junction knockdown | 3rd | R65C12-GAL4.DBD / UAS-shakB[RNAi] (VALIUM20) | BDSC #68356 (OL0005B split-GAL4) / BDSC #57706 |
| 4a | LC17 | Co-localzed LC17 and shakB-Tojan-Gal4 | 1st | MiMiC15228-shakB-Trojan-Gal4 / 10xUAS-myr::smGdP-HA, 13xLexAop-myr::smGdP-V5 | A. Borst Lab / G. Rubin lab, Janelia |
| | | | 2nd | R21D03-LexA / + | BDSC #52590 |
| | | | 3rd | + / + | |
| 4a | T3/T2a | Co-localzed T3/T2a and shakB-Tojan-Gal4 | 1st | MiMiC15228-shakB-Trojan-Gal4 / 10xUAS-myr::smGdP-HA, 13xLexAop-myr::smGdP-V5 | A. Borst Lab / G. Rubin lab, Janelia |
| | | | 2nd | R65B04-LexA / + | BDSC #61579 |
| | | | 3rd | + / + | |
| 4d | LC17 | Expressing nuclear GFP | 1st | w[1118] / w[1118] | |
| | | HCR-FISH | 2nd | R21D03-p65.AD / UAS-His2A-GFP | L. Zipursky Lab |
| | | | 3rd | R65C12-GAL4.DBD / + | BDSC #68356 (OL0005B split-GAL4) |
| 4d | T3 | Expressing nuclear GFP | 1st | w[1118] / w[1118] | |
| | | HCR-FISH | 2nd | VT002055-p65.AD / UAS-His2A-GFP | BDSC #73391 / L. Zipursky Lab |
| | | | 3rd | R65B04-GAL4.DBD / + | BDSC #69453 (Keleş et al., 2020) |
| 4d | T2a | Expressing nuclear GFP | 1st | w[1118] / w[1118] | |
| | | HCR-FISH | 2nd | VT012791-p65.AD / UAS-His2A-GFP | BDSC #74047 / L. Zipursky Lab |
| | | | 3rd | R47E02-GAL4.DBD / + | BDSC #69376 (Keleş et al., 2020) |
| 4d | LC11 | Expressing nuclear GFP | 1st | w[1118] / w[1118] | |
| Ext. Fig. 6b | | HCR-FISH | 2nd | R22H02-p65.AD / UAS-His2A-GFP | L. Zipursky Lab |
| | | | 3rd | R20G06-GAL4.DBD / + | BDSC #68362 (OL0015B split-GAL4) |
| 4d | LC16 | Expressing nuclear GFP | 1st | w[1118] / w[1118] | |
| Ext. Fig. 6b | | HCR-FISH | 2nd | R28D11-p65.AD / UAS-His2A-GFP | L. Zipursky Lab |
| | | | 3rd | R54A05-GAL4.DBD / + | BDSC #68360 (OL0017B split-GAL4) |
| 4d | LC12 | Expressing nuclear GFP | 1st | w[1118] / w[1118] | |
| Ext. Fig. 6b | | HCR-FISH | 2nd | R35D04-p65.AD / UAS-His2A-GFP | L. Zipursky Lab |
| | | | 3rd | R55F01-GAL4.DBD / + | BDSC #68360 (OL0008B split-GAL4) |
| 4f,g | LC17 | Co-localzed LC17 and shakB-V5 | 1st | w[1118] / shakB-smGdP-10xV5 | L. Zipursky Lab |
| | | | 2nd | R21D03-p65.AD / 10xUAS-IVS-myr::tdTomato | BDSC #32222 |
| | | | 3rd | R65C12-GAL4.DBD / + | BDSC #68356 (OL0005B split-GAL4) |
| 4g | T3 | Co-localzed T3 and shakB-V5 | 1st | w[1118] / shakB-smGdP-10xV5 | L. Zipursky Lab |
| | | | 2nd | VT002055-p65.AD / 10xUAS-IVS-myr::tdTomato | BDSC #73391 / BDSC #32222 |
| | | | 3rd | R65B04-GAL4.DBD / + | BDSC #69453 (Keleş et al., 2020) |
| 4g | T2a | Co-localzed T2a and shakB-V5 | 1st | w[1118] / shakB-smGdP-10xV5 | L. Zipursky Lab |
| | | | 2nd | VT012791-p65.AD / 10xUAS-IVS-myr::tdTomato | BDSC #74047 / BDSC #32222 |
| | | | 3rd | R47E02-GAL4.DBD / + | BDSC #69376 (Keleş et al., 2020) |
| Ext. Fig. 6c | LC11 | Co-localzed LC11 and shakB-V5 | 1st | w[1118] / shakB-smGdP-10xV5 | L. Zipursky Lab |
| | | | 2nd | R22H02-p65.AD / 10xUAS-IVS-myr::tdTomato | BDSC #32222 |
| | | | 3rd | R20G06-GAL4.DBD / + | BDSC #68362 (OL0015B split-GAL4) |
| Ext. Fig. 6c | LC16 | Co-localzed LC16 and shakB-V5 | 1st | w[1118] / shakB-smGdP-10xV5 | L. Zipursky Lab |

| | | | | | |
|---|---|---|---|---|---|
| | | | 2nd | R28D11-p65.AD / 10xUAS-IVS-myr::tdTomato | BDSC #32222 |
| | | | 3rd | R54A05-GAL4.DBD / + | BDSC #68360 (OL0017B split-GAL4) |
| Ext. Fig. 6c | LC12 | Co-localzed LC12 and shakB-V5 | 1st | w[1118] / shakB-smGdP-10xV5 | L. Zipursky Lab |
| | | | 2nd | R35D04-p65.AD / 10xUAS-IVS-myr::tdTomato | BDSC #32222 |
| | | | 3rd | R55F01-GAL4.DBD / + | BDSC #68360 (OL0008B split-GAL4) |
| 4i | LC17 | Conditional shakB-V5 tagging in LC17 | 1st | w[1118] / shakB-KDRT-STOP-KDRT-smGdP-10xV5 | L. Zipursky Lab |
| | | | 2nd | R21D03-p65.AD / 10xUAS-IVS-myr::GFP-2A-KDR | L. Zipursky Lab |
| | | | 3rd | R65C12-GAL4.DBD / + | BDSC #68356 (OL0005B split-GAL4) |
| 4j | T2a | Conditional shakB-V5 tagging in T2a | 1st | w[1118] / shakB-KDRT-STOP-KDRT-smGdP-10xV5 | L. Zipursky Lab |
| | | | 2nd | VT013616-p65.AD / 10xUAS-IVS-myr::GFP-2A-KDR | L. Zipursky Lab |
| | | | 3rd | VT025776-GAL4.DBD / + | BDSC #88945 (SS02353 split-GAL4) |
| 4j | T3 | Conditional shakB-V5 tagging in T3 | 1st | w[1118] / shakB-KDRT-STOP-KDRT-smGdP-10xV5 | L. Zipursky Lab |
| Ext. Fig. 7 | | | 2nd | 10xUAS-IVS-myr::GFP-2A-KDR / + | L. Zipursky Lab |
| | | | 3rd | VT002055-Gal4 / + | B. Dickson Lab (Keleş et al., 2020) |
| Ext. Fig. 6d | LC4 | Conditional shakB-V5 tagging in LC4 | 1st | w[1118] / shakB-KDRT-STOP-KDRT-smGdP-10xV5 | L. Zipursky Lab |
| | | | 2nd | R47H03-p65.AD / 10xUAS-IVS-myr::GFP-2A-KDR | L. Zipursky Lab |
| | | | 3rd | R72E01-GAL4.DBD / + | BDSC #68259 (SS00315 split-GAL4) |
| Ext. Fig. 6e | T4/T5 | Conditional shakB-V5 tagging in T4/T5 | 1st | w[1118] / shakB-KDRT-STOP-KDRT-smGdP-10xV5 | L. Zipursky Lab |
| | | | 2nd | R59E08-p65.AD / 10xUAS-IVS-myr::GFP-2A-KDR | L. Zipursky Lab |
| | | | 3rd | R42F06-GAL4.DBD / + | BDSC #86858 (SS00324 split-GAL4) |
| 4k | LC17 | Neurobiotin filling in LC17 | 1st | w[1118] / w[1118] | |
| Ext. Fig. 6f | | | 2nd | R21D03-p65.AD / 10xUAS-IVS-mCD8::GFP | BDSC #32188 |
| | | | 3rd | R65C12-GAL4.DBD / UAS-Fmr1[RNAi] (VALIUM20) | BDSC #68356 (OL0005B split-GAL4) / BDSC #34944 |
| Ext. Fig. 7 | T3 | Conditional shakB-V5 tagging in T3 | 1st | w[1118] / shakB-KDRT-STOP-KDRT-smGdP-10xV5 | L. Zipursky Lab |
| | | | 2nd | VT002055-p65.AD / 10xUAS-IVS-myr::GFP-2A-KDR | BDSC #73391 / L. Zipursky Lab |
| | | | 3rd | R65B04-GAL4.DBD / + | BDSC #69453 (Keleş et al., 2020) |
| 5c | Control | Calcium imaging | 1st | 13xLexAop-IVS-jGCaMP7f / w[1118] | BDSC #80910 |
| | | shakB[RNAi] - Gap junction knockdown | 2nd | R21D03-LexA / empty-p65.AD | BDSC #52590 |
| | | | 3rd | empty-GAL4.DBD / UAS-shakB[RNAi] (VALIUM20) | BDSC #79603 (empty split-GAL4) / BDSC #57706 |
| 5c | LC17 | Calcium imaging | 1st | 13xLexAop-IVS-jGCaMP7f / w[1118] | BDSC #80910 |
| | | shakB[RNAi] - Gap junction knockdown | 2nd | R21D03-LexA / VT002055-p65.AD | BDSC #52590 / BDSC #73391 |
| | | | 3rd | R65B04-GAL4.DBD / UAS-shakB[RNAi] (VALIUM20) | BDSC #69453 (Keleş et al., 2020) / BDSC #57706 |
