## Supplemental Table 2 for "Electrical synapses mediate visual approach behavior"

**Figure 1g**
A generalized linear model (GLM) with Gaussian family and identity link function was fitted to the data on calcium response across 3D-SHOT laser powers. A two-way ANOVA (6x2) was then run on the model estimates.

| Predictor | Degrees of Freedom | Residual Deviance | F-value | p-value |
|---|---|---|---|---|
| NULL | 78 | 0.0056 | | |
| Genotype | (1, 76) | 0.0048 | 6.9305 | 0.002 ** |
| Power | (1, 75) | 0.0046 | 3.5984 | 0.06 |
| Genotype*Power | (1, 74) | 0.0043 | 6.0433 | 0.02 * |

**Figure 1i**
* $p < 0.05$; ** $p < 0.01$; and *** $p < 0.001$
One-way ANOVA ($F_{(5, 25)} = 11.56$, $p < 0.0001$) run on calcium responses to different holographic stimulations followed by pairwise t-test comparisons with Holm-Bonferroni correction.

| Comparison | Difference between means ± SE | t-ratio | p-value | Cohen's d |
|---|---|---|---|---|
| S1 – S2 | -0.00114 ± 0.00557 | -0.204 | 1.00 | -0.12 |
| S1 – S3 | 0.00532 ± 0.00557 | 0.955 | 1.00 | 0.55 |
| S1 – S1-3 | -0.00808 ± 0.00557 | -1.452 | 0.95 | -0.84 |
| S1 – Virtual Sum | -0.01272 ± 0.00557 | -2.286 | 0.25 | -1.32 |
| S2 – S3 | 0.00645 ± 0.00557 | 1.159 | 1.00 | 0.67 |
| S2 – S1-3 | -0.00695 ± 0.00557 | -1.248 | 1.00 | -0.72 |
| S2 – Virtual Sum | -0.01159 ± 0.00557 | -2.082 | 0.33 | -1.20 |
| S3 – S1-3 | -0.01340 ± 0.00557 | -2.408 | 0.21 | -1.39 |
| S3 – Virtual Sum | -0.01804 ± 0.00557 | -3.241 | 0.03 * | -1.87 |
| S1-3 – Virtual Sum | -0.00464 ± 0.00557 | -0.833 | 1.00 | -0.48 |

**Figure 2c**
A LM was fitted to the data on peak calcium responses to a moving bar in LC17 and LC11 neurons. An unpaired t-test was then run on the model estimates.

| Comparison | estimate | SE | t-ratio | p-value | Cohen's d |
|---|---|---|---|---|---|
| LC17 – LC11 | 0.0376 | 0.0122 | 3.074 | 0.01 | 1.77 |

**Figure 2g**
A GLM with Gaussian family and identity link function was fitted to the data on calcium response across technical replicates. A two-way ANOVA (3x2) was then run on the model estimates. ON bars moving front-to-back in LC17 and LC11 neurons.

| Predictor | Degrees of Freedom | Residual Deviance | F-value | p-value |
|---|---|---|---|---|
| NULL | 36 | 0.0250 | | |
| Genotype | (2, 34) | 0.0166 | 9.6333 | 0.0006 *** |
| Replicate | (2, 32) | 0.0153 | 1.5220 | 0.23 |
| Genotype*Replicate | (2, 30) | 0.0131 | 2.4570 | 0.10 |

ON bars moving back-to-front in LC17 and LC11 neurons.

| Predictor | Degrees of Freedom | Residual Deviance | F-value | p-value |
|---|---|---|---|---|
| NULL | 36 | 0.0291 | | |
| Genotype | (2, 34) | 0.013021 | 29.3794 | < 0.0001 *** |
| Replicate | (2, 32) | 0.010241 | 5.0762 | 0.01 * |
| Genotype*Replicate | (2, 30) | 0.008214 | 3.7016 | 0.04 * |

**Figure 2i**
A linear mixed-effect model (LMM) was fitted to the data on calcium response. An unpaired t-test was then run on the model fixed-effects estimation. Comparison of peak calcium responses to a dark looming in LC17 and LPLC2 neurons.

| Comparison | estimate | SE | t-ratio | p-value | Cohen's d |
|---|---|---|---|---|---|
| LC17 – LPLC2 | -0.0541 | 0.0175 | -3.086 | 0.005 ** | -1.18 |

**Extended Data Figure 3b**
A LMM was fitted to the data on calcium response. An unpaired t-test was then run on the model fixed-effects estimation. Comparison of peak calcium responses to ON moving bars in LC17 and LC11 neurons.

| Comparison | estimate | SE | t-ratio | p-value | Cohen's d |
|---|---|---|---|---|---|
| LC17 – LC11 | 0.0376 | 0.0101 | 3.741 | 0.002 ** | 1.53 |

Comparison of peak calcium responses to OFF moving bars in LC17 and LC11 neurons.

| Comparison | estimate | SE | t-ratio | p-value | Cohen's d |
|---|---|---|---|---|---|
| LC17 – LC11 | 0.0143 | 0.0067 | 2.123 | 0.051 | 0.87 |

**Extended Data Figure 3c**
A GLM with Gaussian family and identity link function was fitted to the data on calcium response across technical replicates. A two-way ANOVA (3x2) was then run on the model estimates. OFF bars moving front-to-back in LC17 and LC11 neurons.

| Predictor | Degrees of Freedom | Residual Deviance | F-value | p-value |
|---|---|---|---|---|
| NULL | 36 | 0.0200 | | |
| Genotype | (2, 34) | 0.0133 | 8.6989 | 0.001 ** |
| Replicate | (2, 32) | 0.0126 | 0.9224 | 0.41 |
| Genotype*Replicate | (2, 30) | 0.0116 | 1.3397 | 0.28 |

OFF bars moving back-to-front.

| Predictor | Degrees of Freedom | Residual Deviance | F-value | p-value |
|---|---|---|---|---|
| NULL | 35 | 0.0158473 | | |
| Genotype | (2, 33) | 0.0073014 | 18.9282 | < 0.0001 *** |
| Replicate | (2, 31) | 0.0069611 | 0.7537 | 0.48 |
| Genotype*Replicate | (2, 29) | 0.0065466 | 0.9181 | 0.41 |

**Figure 3d**
A GLM with Gamma family and log link function was fitted to the data on performance index values across genotypes. Planned pairwise t-test comparisons were then run on the model estimates. Driver lines crossed with UAS-Kir2.1.

| Comparison | Difference between means ± SE | t-ratio | p-value | Cohen's d |
|---|---|---|---|---|
| EmptySp>Kir2.1 – LC17Sp>Kir2.1 | 0.6981 ± 0.278 | 2.514 | 0.01 * | 0.79 |
| EmptySp>Kir2.1 – T3Sp>Kir2.1 | 0.8584 ± 0.281 | 3.051 | 0.003 ** | 0.98 |
| EmptySp>Kir2.1 – T4T5Sp>Kir2.1 | 0.0362 ± 0.271 | 0.133 | 0.89 | 0.04 |
| EmptySp>Kir2.1 – T2aSp>Kir2.1 | 0.0729 ± 0.306 | 0.238 | 0.81 | 0.08 |

**Figure 3e**
A LM was fitted to the data on frequency of saccade values across genotypes. Planned pairwise t-test comparisons were then run on the model estimates. Driver lines crossed with UAS-Kir2.1.

| Comparison | Difference between means ± SE | t-ratio | p-value | Cohen's d |
|---|---|---|---|---|
| EmptySp>Kir2.1 – LC17Sp>Kir2.1 | 0.3804 ± 0.0713 | 5.340 | < 0.0001 *** | 0.99 |
| EmptySp>Kir2.1 – T3Sp>Kir2.1 | 0.3423 ± 0.0695 | 4.926 | < 0.0001 *** | 0.89 |
| EmptySp>Kir2.1 – T4T5Sp>Kir2.1 | 0.4376 ± 0.0748 | 5.851 | < 0.0001 *** | 1.14 |
| EmptySp>Kir2.1 – T2aSp>Kir2.1 | 0.2080 ± 0.0688 | 3.021 | 0.003 ** | 0.54 |

**Figure 3g**
A GLM with Gamma family and log link function was fitted to the data on performance index values across genotypes. Planned pairwise t-test comparisons were then run on the model estimates. Driver lines crossed with UAS-TeTxLC on X.

| Comparison | Difference between means ± SE | t-ratio | p-value | Cohen's d |
|---|---|---|---|---|
| EmptySp>TeTxLC(X) – LC17Sp>TeTxLC(X) | -0.4688 ± 0.304 | -1.542 | 0.13 | -0.41 |
| EmptySp>TeTxLC(X) – T3Sp>TeTxLC(X) | -0.4274 ± 0.304 | -1.406 | 0.16 | -0.37 |
| EmptySp>TeTxLC(X) – T4T5Sp>TeTxLC(X) | -2.6869 ± 0.388 | -6.933 | < 0.0001 *** | -2.33 |
| EmptySp>TeTxLC(X) – T2aSp>TeTxLC(X) | -0.6943 ± 0.320 | -2.168 | 0.03 * | -0.60 |

**Figure 3h**
A LM was fitted to the data on frequency of saccade values across genotypes. Planned pairwise t-test comparisons were then run on the model estimates. Driver lines crossed with UAS-TeTxLC on X.

| Comparison | Difference between means ± SE | t-ratio | p-value | Cohen's d |
|---|---|---|---|---|
| EmptySp>TeTxLC(X) – LC17Sp>TeTxLC(X) | 0.142330 ± 0.0421 | 3.378 | 0.0008 *** | 0.48 |
| EmptySp>TeTxLC(X) – T3Sp>TeTxLC(X) | 0.143111 ± 0.0423 | 3.386 | 0.0008 *** | 0.48 |
| EmptySp>TeTxLC(X) – T4T5Sp>TeTxLC(X) | 0.327385 ± 0.0872 | 3.754 | 0.0002 *** | 1.10 |
| EmptySp>TeTxLC(X) – T2aSp>TeTxLC(X) | 0.180732 ± 0.0453 | 3.992 | 0.0001 *** | 0.61 |

**Figure 3j**
A GLM with Gamma family and log link function was fitted to the data on performance index values across genotypes. Planned pairwise t-test comparisons were then run on the model estimates. Driver lines crossed with UAS-ShakB[RNAi].

| Comparison | Difference between means ± SE | t-ratio | p-value | Cohen's d |
|---|---|---|---|---|
| EmptySp>ShakB[RNAi] – LC17Sp>ShakB[RNAi] | 0.587 ± 0.182 | 3.232 | 0.002 ** | 0.91 |
| EmptySp>ShakB[RNAi] – T3Sp>ShakB[RNAi] | 0.339 ± 0.182 | 1.870 | 0.06 | 0.53 |
| EmptySp>ShakB[RNAi] – T4T5Sp>ShakB[RNAi] | -0.239 ± 0.185 | -1.289 | 0.20 | -0.37 |
| EmptySp>ShakB[RNAi] – T2aSp>ShakB[RNAi] | 0.110 ± 0.193 | 0.569 | 0.57 | 0.17 |

**Figure 3k**
A LM was fitted to the data on frequency of saccade values across genotypes. Planned pairwise t-test comparisons were then run on the model estimates. Driver lines crossed with UAS-ShakB[RNAi].

| Comparison | Difference between means ± SE | t-ratio | p-value | Cohen's d |
|---|---|---|---|---|
| EmptySp>ShakB[RNAi] – LC17Sp>ShakB[RNAi] | 0.1526 ± 0.0654 | 2.335 | 0.02 * | 0.33 |
| EmptySp>ShakB[RNAi] – T3Sp>ShakB[RNAi] | 0.1637 ± 0.0655 | 2.498 | 0.01 * | 0.36 |
| EmptySp>ShakB[RNAi] – T4T5Sp>ShakB[RNAi] | -0.1820 ± 0.0678 | -2.686 | 0.008 ** | -0.40 |
| EmptySp>ShakB[RNAi] – T2aSp>ShakB[RNAi] | 0.0343 ± 0.0694 | 0.495 | 0.62 | 0.07 |

**Extended Data Figure 4b**
A GLM with Gamma family and log link function was fitted to the data on performance index values for wide-field panorama across genotypes. Planned pairwise t-test comparisons were then run on the model estimates. Driver lines crossed with UAS-Kir2.1.

| Comparison | Difference between means ± SE | t-ratio | p-value | Cohen's d |
|---|---|---|---|---|
| EmptySp>Kir2.1 – LC17Sp>Kir2.1 | 0.0651 ± 0.0617 | 1.056 | 0.29 | 0.32 |
| EmptySp>Kir2.1 – T3Sp>Kir2.1 | 0.1226 ± 0.0609 | 2.013 | 0.05 | 0.60 |
| EmptySp>Kir2.1 – T4T5Sp>Kir2.1 | 0.4340 ± 0.0609 | 7.123 | < 0.0001 *** | 2.12 |
| EmptySp>Kir2.1 – T2aSp>Kir2.1 | 0.0241 ± 0.0678 | 0.355 | 0.72 | 0.12 |

A LM was fitted to the data on frequency of saccade values during wide-field panorama presentation across genotypes. Planned pairwise t-test comparisons were then run on the model estimates. Driver lines crossed with UAS-Kir2.1.

| Comparison | Difference between means ± SE | t-ratio | p-value | Cohen's d |
|---|---|---|---|---|
| EmptySp>Kir2.1 – LC17Sp>Kir2.1 | -0.1372 ± 0.0557 | -2.465 | 0.01 * | -0.46 |
| EmptySp>Kir2.1 – T3Sp>Kir2.1 | -0.0329 ± 0.0548 | -0.601 | 0.55 | -0.11 |
| EmptySp>Kir2.1 – T4T5Sp>Kir2.1 | 0.0766 ± 0.0588 | 1.303 | 0.19 | 0.26 |
| EmptySp>Kir2.1 – T2aSp>Kir2.1 | 0.1226 ± 0.0541 | 2.268 | 0.02 * | 0.41 |

**Extended Data Figure 4c**
A GLM with Gamma family and log link function was fitted to the data on performance index values for wide-field panorama across genotypes. Planned pairwise t-test comparisons were then run on the model estimates. Driver lines crossed with UAS-TeTxLC on X.

| Comparison | Difference between means ± SE | t-ratio | p-value | Cohen's d |
|---|---|---|---|---|
| EmptySp>TeTxLC(X) – LC17Sp>TeTxLC(X) | -0.0944 ± 0.190 | -0.496 | 0.62 | -0.13 |
| EmptySp>TeTxLC(X) – T3Sp>TeTxLC(X) | -0.0162 ± 0.186 | -0.087 | 0.93 | -0.02 |
| EmptySp>TeTxLC(X) – T4T5Sp>TeTxLC(X) | -0.7937 ± 0.255 | -3.108 | 0.002 ** | -1.07 |
| EmptySp>TeTxLC(X) – T2aSp>TeTxLC(X) | 0.0379 ± 0.200 | 0.189 | 0.85 | 0.05 |

A LM was fitted to the data on frequency of saccade values during wide-field panorama presentation across genotypes. Planned pairwise t-test comparisons were then run on the model estimates. Driver lines crossed with UAS-TeTxLC on X.

| Comparison | Difference between means ± SE | t-ratio | p-value | Cohen's d |
|---|---|---|---|---|
| EmptySp>TeTxLC(X) – LC17Sp>TeTxLC(X) | 0.0280 ± 0.0348 | 0.803 | 0.42 | 0.12 |
| EmptySp>TeTxLC(X) – T3Sp>TeTxLC(X) | -0.0000993 ± 0.0352 | -0.003 | 0.10 | -0.0004 |
| EmptySp>TeTxLC(X) – T4T5Sp>TeTxLC(X) | 0.0257 ± 0.0731 | 0.352 | 0.72 | 0.11 |
| EmptySp>TeTxLC(X) – T2aSp>TeTxLC(X) | 0.0404 ± 0.0374 | 1.080 | 0.28 | 0.18 |

**Extended Data Figure 4d**

A GLM with Gamma family and log link function was fitted to the data on performance index values for wide-field panorama across genotypes. Planned pairwise t-test comparisons were then run on the model estimates. Driver lines crossed with UAS-ShakB[RNAi].

| Comparison | Difference between means ± SE | t-ratio | p-value | Cohen's d |
|---|---|---|---|---|
| EmptySp>ShakB[RNAi] – LC17Sp>ShakB[RNAi] | -0.025270 ± 0.0477 | -0.530 | 0.60 | -0.15 |
| EmptySp>ShakB[RNAi] – T3Sp>ShakB[RNAi] | 0.047084 ± 0.0477 | 0.988 | 0.33 | 0.28 |
| EmptySp>ShakB[RNAi] – T4T5Sp>ShakB[RNAi] | 0.162988 ± 0.0487 | 3.348 | 0.001 ** | 0.97 |
| EmptySp>ShakB[RNAi] – T2aSp>ShakB[RNAi] | -0.022859 ± 0.0505 | -0.452 | 0.65 | -0.14 |

A LM was fitted to the data on frequency of saccade values during wide-field panorama presentation across genotypes. Planned pairwise t-test comparisons were then run on the model estimates. Driver lines crossed with UAS-ShakB[RNAi].

| Comparison | Difference between means ± SE | t-ratio | p-value | Cohen's d |
|---|---|---|---|---|
| EmptySp>ShakB[RNAi] – LC17Sp>ShakB[RNAi] | 0.02034 ± 0.0324 | 0.627 | 0.53 | 0.09 |
| EmptySp>ShakB[RNAi] – T3Sp>ShakB[RNAi] | 0.15156 ± 0.0337 | 4.497 | < 0.0001 *** | 0.70 |
| EmptySp>ShakB[RNAi] – T4T5Sp>ShakB[RNAi] | 0.08520 ± 0.0331 | 2.572 | 0.01 * | 0.39 |
| EmptySp>ShakB[RNAi] – T2aSp>ShakB[RNAi] | 0.14073 ± 0.0354 | 3.977 | 0.0001 *** | 0.65 |

**Extended Data Figure 4f**
A GLM with Gamma family and log link function was fitted to the data on performance index values for small-object across genotypes. Planned pairwise t-test comparisons were then run on the model estimates. Driver lines crossed with UAS-TeTxLC on X.

| Comparison | Difference between means ± SE | t-ratio | p-value | Cohen's d |
|---|---|---|---|---|
| EmptySp>TeTxLC(X) – LC17Sp>TeTxLC(X) | -0.672 ± 0.580 | -1.158 | 0.25 | -0.40 |
| EmptySp>TeTxLC(X) – T3Sp>TeTxLC(X) | -0.765 ± 0.604 | -1.267 | 0.21 | -0.46 |
| EmptySp>TeTxLC(X) – T2aSp>TeTxLC(X) | -1.060 ± 0.563 | -1.884 | 0.06 | -0.63 |

A LM was fitted to the data on frequency of saccade values during small-object presentation across genotypes. Planned pairwise t-test comparisons were then run on the model estimates. Driver lines crossed with UAS-TeTxLC on X.

| Comparison | Difference between means ± SE | t-ratio | p-value | Cohen's d |
|---|---|---|---|---|
| EmptySp>TeTxLC(X) – LC17Sp>TeTxLC(X) | 0.0787 ± 0.0419 | 1.880 | 0.06 | 0.34 |
| EmptySp>TeTxLC(X) – T3Sp>TeTxLC(X) | 0.0955 ± 0.0442 | 2.160 | 0.03 * | 0.41 |
| EmptySp>TeTxLC(X) – T2aSp>TeTxLC(X) | 0.1552 ± 0.0417 | 3.720 | 0.0002 *** | 0.67 |

A GLM with Gamma family and log link function was fitted to the data on performance index values for small-object across genotypes. Planned pairwise t-test comparisons were then run on the model estimates. Driver lines crossed with UAS-ShakB[RNAi].

| Comparison | Difference between means ± SE | t-ratio | p-value | Cohen's d |
|---|---|---|---|---|
| EmptySp>ShakB[RNAi] – LC17Sp>ShakB[RNAi] | 0.3257 ± 0.254 | 1.284 | 0.20 | 0.36 |
| EmptySp>ShakB[RNAi] – T3Sp>ShakB[RNAi] | 0.3992 ± 0.256 | 1.557 | 0.12 | 0.45 |
| EmptySp>ShakB[RNAi] – T4T5Sp>ShakB[RNAi] | 0.6615 ± 0.259 | 2.550 | 0.01 * | 0.75 |
| EmptySp>ShakB[RNAi] – T2aSp>ShakB[RNAi] | 0.9523 ± 0.274 | 3.471 | 0.0007 *** | 1.07 |

A LM was fitted to the data on frequency of saccade values during small-object presentation across genotypes. Planned pairwise t-test comparisons were then run on the model estimates. Driver lines crossed with UAS-ShakB[RNAi].

| Comparison | Difference between means ± SE | t-ratio | p-value | Cohen's d |
|---|---|---|---|---|
| EmptySp>ShakB[RNAi] – LC17Sp>ShakB[RNAi] | 0.06694 ± 0.0332 | 2.016 | 0.04 * | 0.29 |
| EmptySp>ShakB[RNAi] – T3Sp>ShakB[RNAi] | 0.04816 ± 0.0332 | 1.450 | 0.15 | 0.21 |
| EmptySp>ShakB[RNAi] – T4T5Sp>ShakB[RNAi] | -0.01089 ± 0.0338 | -0.322 | 0.75 | -0.05 |
| EmptySp>ShakB[RNAi] – T2aSp>ShakB[RNAi] | 0.12356 ± 0.0350 | 3.528 | 0.0005 *** | 0.53 |

**Extended Data Figure 4g**

A GLM with Gamma family and log link function was fitted to the data on performance index values for motion-defined bar across genotypes. Planned pairwise t-test comparisons were then run on the model estimates. Driver lines crossed with UAS-TeTxLC on II.

| Comparison | Difference between means ± SE | t-ratio | p-value | Cohen's d |
|---|---|---|---|---|
| EmptySp>TeTxLC(II) – LC17Sp>TeTxLC(II) | -0.3714 ± 0.282 | -1.318 | 0.19 | -0.40 |
| EmptySp>TeTxLC(II) – T3Sp>TeTxLC(II) | 0.0796 ± 0.289 | 0.275 | 0.78 | 0.09 |
| EmptySp>TeTxLC(II) – T4T5Sp>TeTxLC(II) | -2.0296 ± 0.289 | -7.018 | < 0.0001 *** | -2.20 |
| EmptySp>TeTxLC(II) – T2aSp>TeTxLC(II) | -0.2798 ± 0.282 | -0.993 | 0.32 | -0.30 |

A LM was fitted to the data on frequency of saccade values during motion-defined presentation across genotypes. Planned pairwise t-test comparisons were then run on the model estimates. Driver lines crossed with UAS-TeTxLC on II.

| Comparison | Difference between means ± SE | t-ratio | p-value | Cohen's d |
|---|---|---|---|---|
| EmptySp>TeTxLC(II) – LC17Sp>TeTxLC(II) | -0.05373 ± 0.0467 | -1.151 | 0.25 | -0.18 |
| EmptySp>TeTxLC(II) – T3Sp>TeTxLC(II) | 0.03484 ± 0.0467 | 0.746 | 0.46 | 0.11 |
| EmptySp>TeTxLC(II) – T4T5Sp>TeTxLC(II) | 0.11755 ± 0.0517 | 2.275 | 0.02 * | 0.39 |
| EmptySp>TeTxLC(II) – T2aSp>TeTxLC(II) | -0.05815 ± 0.0468 | -1.242 | 0.22 | -0.19 |

A GLM with Gamma family and log link function was fitted to the data on performance index values for wide-field panorama across genotypes. Planned pairwise t-test comparisons were then run on the model estimates. Driver lines crossed with UAS-TeTxLC on II. The comparison between EmptySp and T4/T5Sp was performed by filtering values greater than or equal to 1 in order to capture better behavioral dynamics.

| Comparison | Difference between means ± SE | t-ratio | p-value | Cohen's d |
|---|---|---|---|---|
| EmptySp>TeTxLC(II) – LC17Sp>TeTxLC(II) | 0.2553 ± 0.181 | 1.413 | 0.16 | 0.43 |
| EmptySp>TeTxLC(II) – T3Sp>TeTxLC(II) | 0.1159 ± 0.183 | 0.634 | 0.53 | 0.19 |
| EmptySp>TeTxLC(II) – T4T5Sp>TeTxLC(II) | -0.84457 ± 0.088 | -9.572 | < 0.0001 *** | -3.84 |
| EmptySp>TeTxLC(II) – T2aSp>TeTxLC(II) | -0.0475 ± 0.183 | -0.260 | 0.80 | -0.08 |

A LM was fitted to the data on frequency of saccade values during wide-field panorama presentation across genotypes. Planned pairwise t-test comparisons were then run on the model estimates. Driver lines crossed with UAS-TeTxLC on II.

| Comparison | Difference between means ± SE | t-ratio | p-value | Cohen's d |
|---|---|---|---|---|
| EmptySp>TeTxLC(II) – LC17Sp>TeTxLC(II) | -0.0807 ± 0.0432 | -1.868 | 0.06 | -0.30 |
| EmptySp>TeTxLC(II) – T3Sp>TeTxLC(II) | 0.0283 ± 0.0458 | 0.617 | 0.54 | 0.11 |
| EmptySp>TeTxLC(II) – T4T5Sp>TeTxLC(II) | 0.1243 ± 0.0498 | 2.499 | 0.01 * | 0.46 |
| EmptySp>TeTxLC(II) – T2aSp>TeTxLC(II) | -0.0418 ± 0.0469 | -0.892 | 0.37 | -0.16 |

A GLM with Gamma family and log link function was fitted to the data on performance index values for small-object across genotypes. Planned pairwise t-test comparisons were then run on the model estimates. Driver lines crossed with UAS-TeTxLC on II.

| Comparison | Difference between means ± SE | t-ratio | p-value | Cohen's d |
|---|---|---|---|---|
| EmptySp>TeTxLC(II) – LC17Sp>TeTxLC(II) | -0.0372 ± 0.268 | -0.139 | 0.89 | -0.04 |
| EmptySp>TeTxLC(II) – T3Sp>TeTxLC(II) | 0.3918 ± 0.271 | 1.447 | 0.15 | 0.45 |
| EmptySp>TeTxLC(II) – T4T5Sp>TeTxLC(II) | -2.0001 ± 0.274 | -7.297 | < 0.0001 *** | -2.28 |
| EmptySp>TeTxLC(II) – T2aSp>TeTxLC(II) | 0.4397 ± 0.271 | 1.624 | 0.11 | 0.50 |

A LM was fitted to the data on frequency of saccade values during small-object presentation across genotypes. Planned pairwise t-test comparisons were then run on the model estimates. Driver lines crossed with UAS-TeTxLC on II.

| Comparison | Difference between means ± SE | t-ratio | p-value | Cohen's d |
|---|---|---|---|---|
| EmptySp>TeTxLC(II) – LC17Sp>TeTxLC(II) | -0.0011 ± 0.0433 | -0.026 | 0.98 | -0.004 |
| EmptySp>TeTxLC(II) – T3Sp>TeTxLC(II) | 0.1177 ± 0.0431 | 2.729 | 0.007 ** | 0.42 |
| EmptySp>TeTxLC(II) – T4T5Sp>TeTxLC(II) | 0.1712 ± 0.0491 | 3.487 | 0.0005 *** | 0.61 |
| EmptySp>TeTxLC(II) – T2aSp>TeTxLC(II) | 0.1064 ± 0.0433 | 2.459 | 0.01 * | 0.38 |

**Extended Data Figure 5b**

A LM was fitted to the data on HCR-FISH puncta in LC17>Control and LC17>UAS-ShakB[RNAi]. An unpaired t-test was then run on the model estimates.

| Comparison | estimate | SE | t-ratio | p-value | Cohen's d |
|---|---|---|---|---|---|
| Ctrl – LC17 | 3.79 | 0.899 | 4.214 | 0.003 ** | 2.67 |

**Figure 4e**

A LM was fitted to the data on HCR-FISH puncta in different visual neurons. Planned pairwise t-test comparisons were then run on the model estimates.

| Comparison | Difference between means ± SE | t-ratio | p-value | Cohen's d |
|---|---|---|---|---|
| LC17 – T3 | 8.659 1.15 | 7.499 | < 0.0001 *** | 4.33 |
| LC17 – LC11 | 6.277 1.15 | 5.436 | < 0.0001 *** | 3.14 |
| LC17 – T2a | 2.882 1.15 | 2.496 | 0.02 * | 1.44 |
| LC17 – LC12 | 2.065 1.15 | 1.788 | 0.08 | 1.03 |
| LC17 – LC16 | 7.321 1.15 | 6.340 | < 0.0001 *** | 3.66 |

**Figure 5c**

Three different models were fitted to the data on peak calcium responses to a moving bar (two different directions) in LC17>Control and LC17>UAS-ShakB[RNAi]. Model comparison was then performed based on the Bayesian Information Criterion (BIC). The BICs from the best two models were finally used to compute an approximation of the Bayes Factor (BF). $BF_{12}$ = 36.97. It means how much more probable is one model than the other.

$$BF_{an} = e^{\frac{(BIC_{null} - BIC_{alternative})}{2}}$$

| Hypothesis | Models | Degrees of Freedom | $R^2$ | AIC | BIC |
|---|---|---|---|---|---|
| 1 | Genotype | 3 | 0.22 | -284.59 | -277.19 |
| 2 | Genotype*Direction | 5 | 0.23 | -282.30 | -269.97 |
| 0 | NULL | 1 | 0 | -267.19 | -264.77 |

**Figure 5d - Extended Data Figure 8**

A GLM with Gamma family and log link function was fitted to the data on peak angular velocity during saccades. An unpaired t-test was then run on the model estimates. Driver lines crossed with UAS-Kir2.1.

| Comparison | estimate | SE | t-ratio | p-value | Cohen's d |
|---|---|---|---|---|---|
| EmptySp – LC17Sp | 0.000269 | 0.0997 | 0.003 | 0.998 | 0.0008 |

Driver lines crossed with UAS-TeTxLC on X.

| Comparison | estimate | SE | t-ratio | p-value | Cohen's d |
|---|---|---|---|---|---|
| EmptySp – LC17Sp | -0.244 | 0.0999 | -2.444 | 0.02 * | -0.63 |

Driver lines crossed with UAS-ShakB[RNAi].

| Comparison | estimate | SE | t-ratio | p-value | Cohen's d |
|---|---|---|---|---|---|
| EmptySp – LC17Sp | 0.274 | 0.0792 | 3.462 | 0.001 * | 0.98 |

**Extended Data Figure 8**

A GLM with Gamma family and identity link function was fitted to the data on the number of tracking bouts. An unpaired t-test was then run on the model estimates. Driver lines crossed with UAS-Kir2.1.

| Comparison | estimate | SE | t-ratio | p-value | Cohen's d |
|---|---|---|---|---|---|
| EmptySp – LC17Sp | 2.33 | 1.06 | 2.204 | 0.03 * | 3.95 |

Driver lines crossed with UAS-TeTxLC on X.

| Comparison | estimate | SE | t-ratio | p-value | Cohen's d |
|---|---|---|---|---|---|
| EmptySp – LC17Sp | 0.0395 | 0.596 | 0.066 | 0.95 | 0.11 |

Driver lines crossed with UAS-ShakB[RNAi].

| Comparison | estimate | SE | t-ratio | p-value | Cohen's d |
|---|---|---|---|---|---|
| EmptySp – LC17Sp | -0.6 | 0.793 | -0.757 | 0.45 | -1.87 |

A LM was fitted to the data on frequency of saccade. An unpaired t-test was then run on the model estimates. Driver lines crossed with UAS-Kir2.1.

| Comparison | estimate | SE | t-ratio | p-value | Cohen's d |
|---|---|---|---|---|---|
| EmptySp – LC17Sp | 0.268 | 0.0689 | 3.883 | 0.0002 *** | 0.76 |

Driver lines crossed with UAS-TeTxLC on X.

| Comparison | estimate | SE | t-ratio | p-value | Cohen's d |
|---|---|---|---|---|---|
| EmptySp – LC17Sp | 0.031 | 0.0451 | 0.688 | 0.49 | 0.11 |

Driver lines crossed with UAS-ShakB[RNAi].

| Comparison | estimate | SE | t-ratio | p-value | Cohen's d |
|---|---|---|---|---|---|
| EmptySp – LC17Sp | 0.132 | 0.0607 | 2.171 | 0.03 * | 0.32 |

A GLM with Gamma family and log link function was fitted to the data on the inter-saccade velocity. An unpaired t-test was then run on the model estimates. Driver lines crossed with UAS-Kir2.1.

| Comparison | estimate | SE | t-ratio | p-value | Cohen's d |
|---|---|---|---|---|---|
| EmptySp – LC17Sp | 0.478 | 0.128 | 3.746 | 0.0005 *** | 1.13 |

Driver lines crossed with UAS-TeTxLC on X.

| Comparison | estimate | SE | t-ratio | p-value | Cohen's d |
|---|---|---|---|---|---|
| EmptySp – LC17Sp | -0.566 | 0.327 | -1.733 | 0.09 | -0.44 |

Driver lines crossed with UAS-ShakB[RNAi].

| Comparison | estimate | SE | t-ratio | p-value | Cohen's d |
|---|---|---|---|---|---|
| EmptySp – LC17Sp | 0.336 | 0.316 | 1.066 | 0.29 | 0.30 |
